# Dynamic Boolean modelling reveals the influence of energy supply on bacterial efflux pump expression

**DOI:** 10.1101/2021.12.13.472361

**Authors:** Ryan Kerr, Sara Jabbari, Jessica M. A. Blair, Iain G. Johnston

**Author notes:** Correspondence to (SJ) or (IGJ).

## Abstract

Antimicrobial resistance (AMR) is a global health issue. One key factor contributing to AMR is the ability of bacteria to export drugs through efflux pumps, which relies on the ATP-dependent expression and interaction of several controlling genes. Recent studies have shown significant cell-to-cell ATP variability exists within clonal bacterial populations, but the contribution of intrinsic cell-to-cell ATP heterogeneity is generally overlooked in understanding efflux pumps. Here, we consider how ATP variability influences gene regulatory networks controlling expression of efflux pump genes in two bacterial species. We develop and apply a generalisable Boolean modelling framework, developed to incorporate the dependence of gene expression dynamics on available cellular energy supply. Theoretical results show differences in energy availability can cause pronounced downstream heterogeneity in efflux gene expression. Cells with higher energy availability have a superior response to stressors. Further, in the absence of stress, model bacteria develop heterogeneous pulses of efflux pump gene expression which contribute to a sustained sub-population of cells with increased efflux expression activity, potentially conferring a continuous pool of intrinsically resistant bacteria. This modelling approach thus reveals an important source of heterogeneity in cell responses to antimicrobials and sheds light on potentially targetable aspects of efflux pump-related antimicrobial resistance.

## 1 Introduction

Antimicrobial resistance (AMR) is a huge global health challenge, contributing to over 700,000 deaths world-wide [1] and estimated to exceed cancer mortality by 2050 if neglected [2]. Protein complexes called efflux pumps are an important mechanism for intrinsic (and acquired) resistance to antibiotics [3–5]. Efflux pumps are capable of transporting a range of substances from the cell compartment to the extracellular medium, reducing intracellular accumulation of substances including antibiotics, and thus conferring resistance. Their central role in facilitating clinically relevant AMR makes efflux pumps an attractive drug target, with much research directed at efflux pump inhibitors as a novel therapeutic to overcome efflux-mediated resistance [6–10].

A current mystery in the activity of efflux pumps is the causes and effects of cell-to-cell heterogeneity in their expression. Recent data demonstrates efflux pump gene expression is not homogeneous throughout the population – instead, bacterial populations display significant cell-to-cell variability [11–13]. Heterogeneity in efflux activity has been suggested to have further downstream influence on AMR. One study, by Sá nchez-Romero and Casadesú s [13], reported a correlation between bacteria with high efflux pump activity in the absence of antibiotic compounds and increased resistance to antibiotics. Additionally, a study by El Meouche and Dunlop [11] has shown a sub-population of bacteria with higher efflux pump expression has lower expression of mismatch repair genes, facilitating mutations which can lead to permanent genetic changes bestowing resistance. However, the underlying dynamics and influences of efflux heterogeneity in cellular populations remain to be uncovered.

One potential source of heterogeneity in gene expression dynamics and interactions is cell-to-cell differences in available energy supply [14, 15]. Heterogeneity in adenosine triphosphate (ATP) levels in bacteria (and other species; see Discussion) is increasingly being characterised experimentally [16, 17]. The quantified range of intracellular ATP can cover almost an order of magnitude, varying between 0.32-2.76 mM in *Escherichia coli* [16], for example. Although not uncontroversial, the ubiquity of cell-to-cell variability in ATP across life suggests it is a genuine effect [18–20]. Following theoretical work showing the potentially profound influence of this variability on gene regulatory networks [14, 15], we here investigate the hypothesis that cell-to-cell differences in available energy can contribute to cell heterogeneity in efflux pump expression, intrinsic resistance to antimicrobials, and response to different environmental stressors.

We previously applied an energy-dependent ordinary differential equation (ODE) framework to a simple GRN, describing the behaviour of many naturally-occurring decision-making circuits [14]. We predicted that differences in cellular energy levels will cause differences in the dynamics and stable outcomes of cellular decision-making. However, while ODE models can in principle be used for GRNs of arbitrary size, the associated parameter space rapidly increases with network size, making it harder to investigate general principles. Here, we develop and use an alternative theoretical framework based on Boolean network models [21, 22], simplifications that have been successfully applied to diverse biological systems [23–25] (although to date not often used to understand bacterial virulence and resistance mechanisms [26, 27]). Despite this widespread and successful use, there has been little consideration of the fact that the processes in Boolean GRN models correspond biologically to ATP-dependent processes [21, 26, 28]. This ATP dependence is important, because it generally influences the dynamics of gene expression [29]. We therefore proceed by developing a simple but highly generalisable modification for including energy dependence in Boolean models of regulatory networks, and use this theoretical framework to explore the effects of ATP variability in experimentally-derived GRN models of efflux pump expression in *E. coli* and *Salmonella*.

## 2 Results

### 2.1 Literature-based model construction accounting for energy availability

We first sought to construct a Boolean modelling framework describing efflux pump expression dynamics, supported by data and yielding verifiable predictions about biological behaviour. We considered AcrAB-TolC, an efflux pump associated with clinically important drug resistance in *E. coli* [30–32] and *Salmonella* [33–36]. TolC is constitutively expressed [37] and predicted to be integrated within at least seven additional efflux transport systems in *E. coli* [38], suggesting the dynamics of *acrAB* are more limiting on producing the complete efflux pump system within each cell. We performed a broad literature review (see Supplementary Information Section 5.1) of efflux pump gene regulation in *E. coli* and *Salmonella* to define a core GRN of *acrAB* in each species (Fig. 1(B)-(C)). This review defined the structure of our model i.e. the nodes, regulatory interactions and edge weights in the corresponding wiring diagrams (Supplementary Fig. S1(A)-(B)).

**Figure 1:**
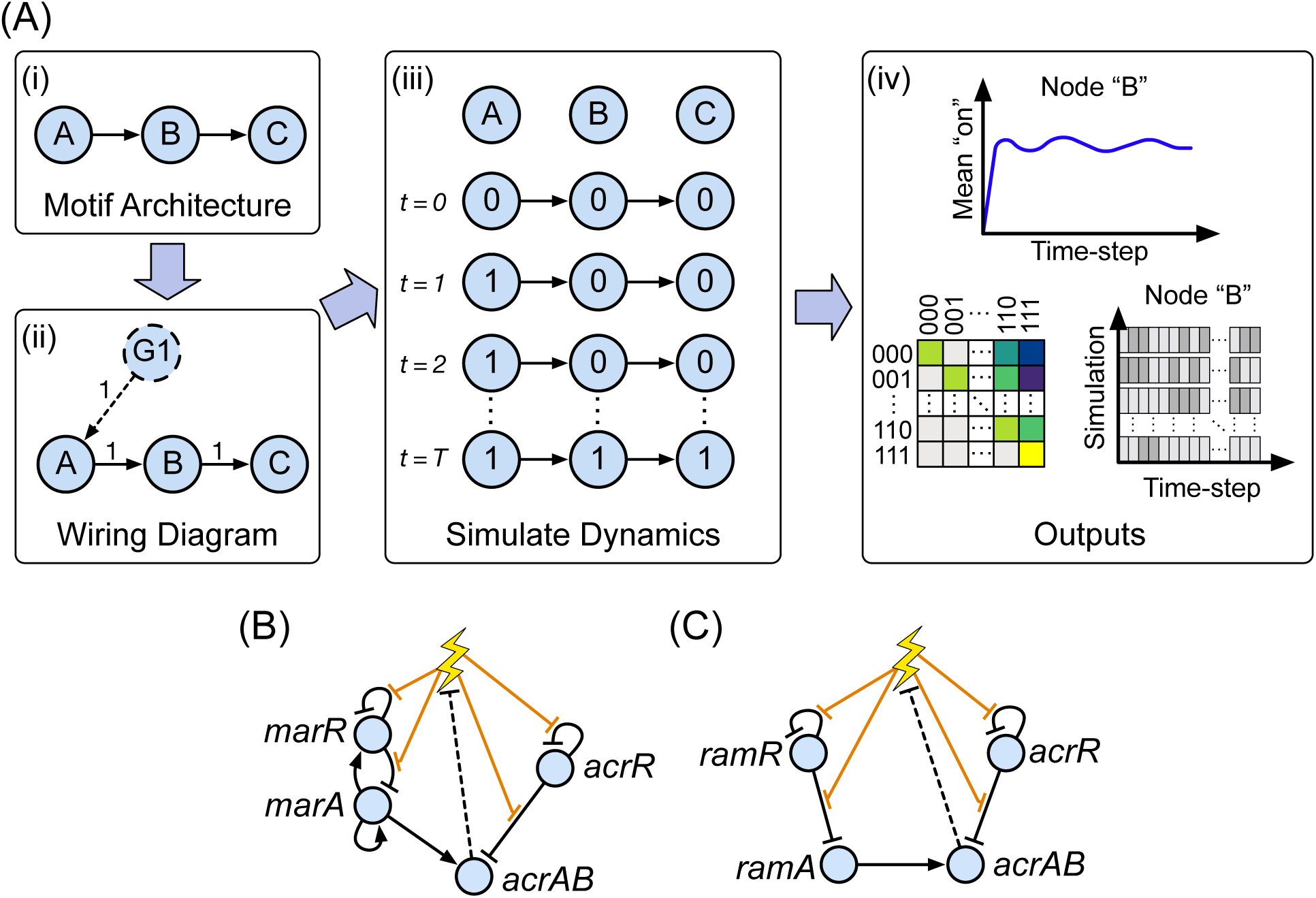
Boolean network modelling of gene regulatory network architectures controlling efflux pump gene expression. **(A)** Schematic overview of the Boolean network framework. Regulatory architectures (i) are converted into wiring diagram form (ii). These wiring diagrams include ‘ghost nodes’ (G1) if necessary, providing a constant signal to ensure constitutively expressed genes are “on” unless switched “off” by a regulator. The considered network is repeatedly simulated over a set number of timesteps (iii), from some initial configuration, producing individual and averaged pictures of network behaviour through transition matrices and expression dynamics of each node per timestep (iv). **(B-C)** Coarse-grained *E. coli* (B) and *Salmonella* (C) AcrAB efflux pump gene regulatory networks, containing a collection of genetic regulators (blue circles) that interact with each other (directed edges) and govern the expression level of efflux genes *acrAB*. Full layouts with parameterizations are given in Supplementary Fig. S1. Antibiotics, or other noxious stressor substances (lightning bolt), can reversibly bind to a gene product (transcription factor) removing its ability to bind to DNA and regulate its target (displayed through orange edges), with consequent downstream effects through the network. Removal of the stressor is integrated through a repressive interaction from *acrAB* to the stress node (dotted edges).

*E. coli* contains a global activator of efflux genes *acrAB*, called *marA* (of the *marRAB* operon), which is repressed locally by the product of *marR* [39]. Similarly, in *Salmonella*, a global activator *ramA* is repressed locally by RamR, the product of *ramR* [40]. In both networks, repressor AcrR, the gene product of *acrR*, down-regulates its own expression and prevents over-expression of efflux genes *acrAB* [41] (see Fig. 1(B)-(C) for each complete GRN). *acrAB* encodes the proteins AcrA and AcrB [42], which can assemble together with TolC to form the AcrAB-TolC system [43, 44] that provides the physical means to expel unwanted substances from the cell (such as toxic metabolic intermediates or bile salts [45, 46]).

The expression of efflux pumps is inducible by the presence of environmental signals, or chemical compounds (such as antibiotics), which we will generally refer to as ‘stressors’. The inducibility of efflux genes enables bacteria to adapt to a wide range of environmental conditions. Noxious substances, such as antibiotics, can bind to GRN products MarR, RamR and AcrR, inhibiting their ability to bind to DNA, and initiating a response cascade that leads to the up-regulation of efflux genes *acrAB* [47–52]. In our model the expression of *acrAB* corresponds to the capacity of the assembled efflux pump to export molecules.

To avoid the aforementioned parameterization issues in continuous models, we developed a general Boolean framework that captures the influence of energy availability. Our model represents genes as nodes in a network, where each gene can be ”on” (expressed) or ”off” (not expressed) at a given time. Edges describe activating and repressing interactions, and the state of a network changes over time with asynchronous update rules following these interactions (see Methods), capturing the stochastic nature of gene regulatory dynamics. To explicitly capture the fact that the fundamental processes (hence all gene interaction ”arrows”) involved in gene expression depend on ATP, we modulate the rate with which regulatory interactions from node *X* to node *Y* are applied according to cellular energy levels (see Methods). This reflects the fact that, for example, cells with low ATP will have a lower rate of transcription elongation per timestep [53], slowing the dynamics of the biochemical intermediates involved in these regulatory interactions, and thus slowing the interactions themselves [14]. While fast fluctuations in ATP supply may exist within cells, as gene expression processes are dynamically slower, we assume the network is exposed to a time-averaged ATP level, and thus define three energy levels ‘high’, ‘intermediate’, and ‘low’, respectively involving a characteristic mean rate of 1, 1*/*2, and 1*/*10 per simulation, corresponding to the aforementioned order of magnitude range observed in biology (see Methods).

### 2.2 Efflux network models reproduce experimentally observed behaviour in *E. coli* and *Salmonella*

Having constructed our model based on a set of experimental observations, we next asked whether it made predictions that could be tested using independent experimental data. In the Supplementary Information we present a table assessing the predictions of our model, for our choice of edge weights, with respect to a multiscale set of experiments at both the population and single-cell level, where agreement is demonstrated across a wide variety of observations (Supplementary Table S1). The overall behaviour of both systems – which we will describe in detail below – predicts a range of coarse- and fine-grained features observed in independent experiments, including pronounced gene expression heterogeneity at the single-cell level, prominent increases in gene expression when a short or long stress signal is provided, the expression of repressive genes in response to stress [41, 54]), and the details of expression levels of different actors in the GRN in the presence and absence of stress.

### 2.3 Stress-free efflux network behaviour with a maximal cellular energy budget

To dissect the dynamics of these networks in detail, we first explored the stress-free population-level behaviour of each network. In the absence of stress, the population-wide mean behaviour in expression levels approaches a steady state (pre-stress dynamics in Fig. 2). We found at high energy availability, initial conditions had no influence on this population-level equilibrium of the *E. coli* and *Salmonella* networks (see Supplementary Information Section 5.5). We therefore proceed by investigating network behaviour starting from a particular initial state – which models an ‘unstressed’ baseline condition – in the rest of the study and focus on the *E. coli* network before comparing the behaviour of the two networks.

**Figure 2:**
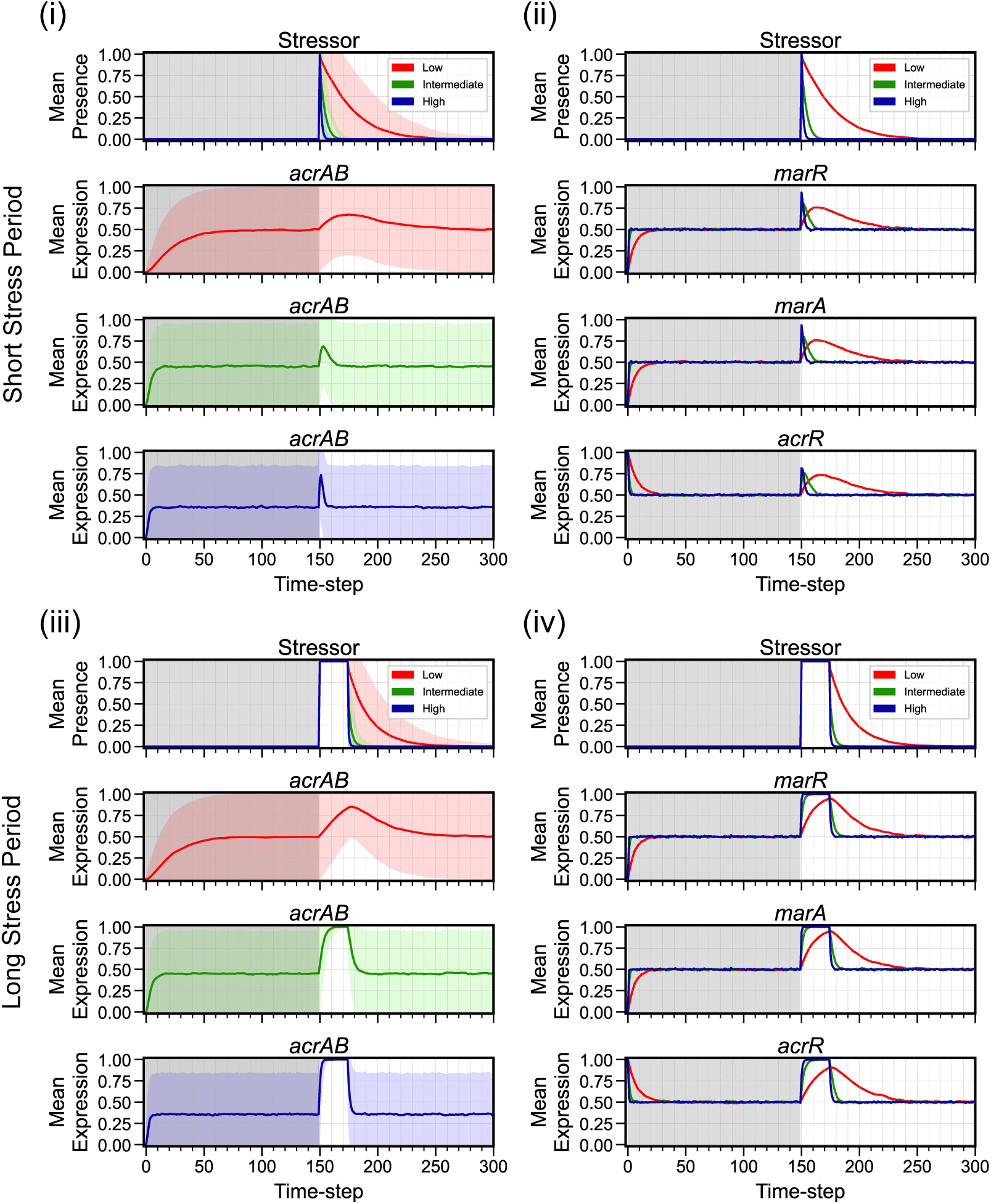
Mean expression dynamics for *E. coli* from the unstressed initial condition in response to a short or long stress period. Time-series dynamics of mean expression (coloured line for low (red), intermediate (green) and high (blue) energy levels) for ((i), (iii)) efflux genes *acrAB* and ((ii), (iv)) remaining regulatory components in *E. coli* ; all panels include stressor behaviour. Behaviour is shown when the network is exposed to a (i)-(ii) short (*t* = 150) or (iii)-(iv) long (*t* = 150-174) stress period, from the unstressed initial condition. The initial time region of each regulatory component prior to stress application (grey shaded region) displays the complete stress-free behaviour. Note: Standard deviation (corresponding coloured shaded area) is included within efflux panels to represent the uncertainty, but the actual expression level cannot exceed the [0, 1] domain.

To explore the variability within the equilibrium population without energetic limitations, we calculated the expression level coefficient of variation (standard deviation divided by mean) for each genetic component in the *E. coli* network. In the absence of stress, expression level coefficient of variation (ELCV), corresponding to population variability in gene expression, ranged between 1.0-1.35 at equilibrium (see stress-free period in Fig. 2). This diversity is due to population-level asynchrony in expression dynamics; in simulated individual bacteria, network components are expressed in heterogeneous pulses (pre-stress dynamics in Fig. 3). This diversity, particularly in *acrAB*, suggests that the network structure itself may support a degree of ‘bet-hedging’ in the absence of stress, where some members of a population are primed to respond to stressors at any given time, agreeing with experimentally observed behaviour in *Salmonella enterica* [13] (see Discussion).

**Figure 3:**
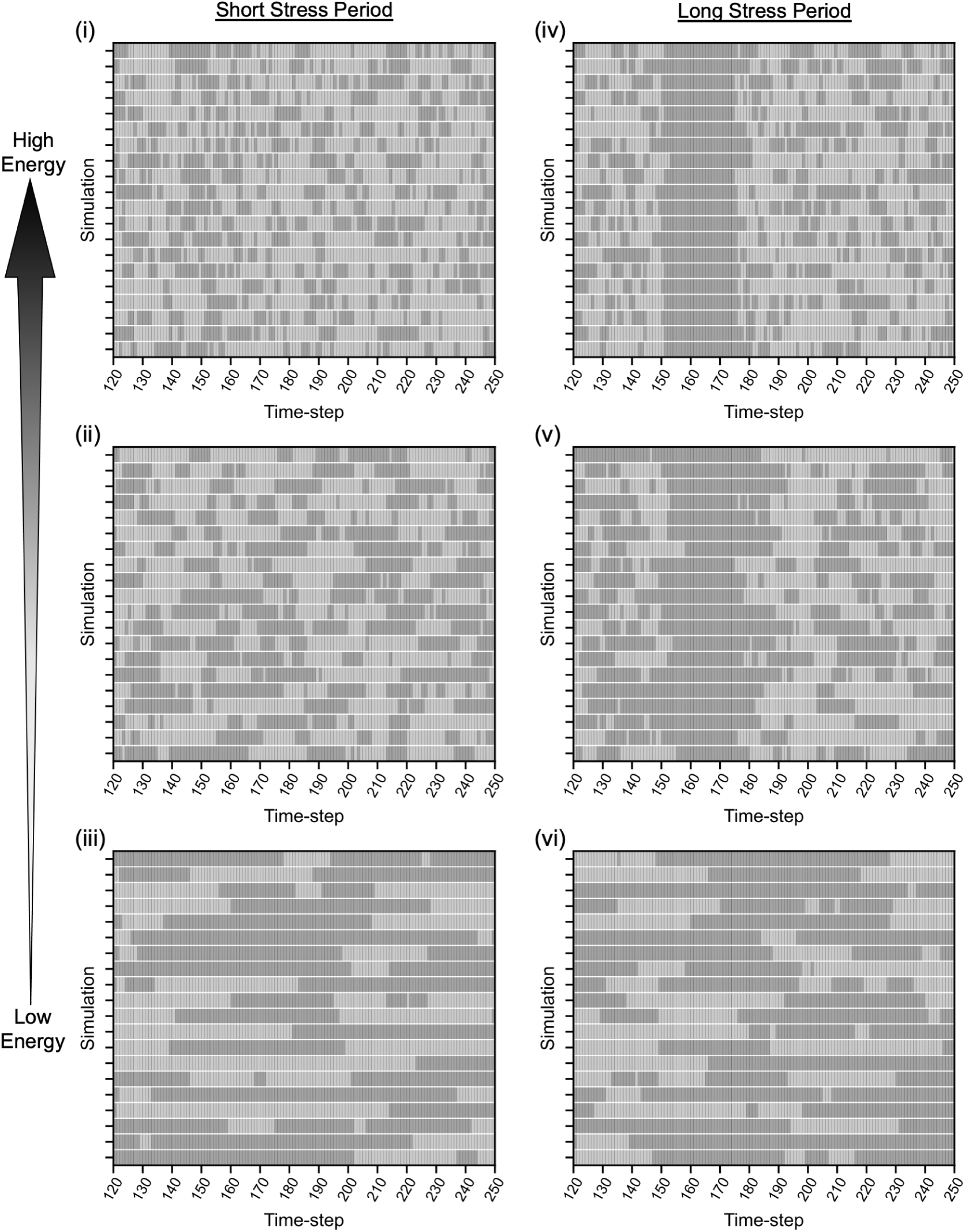
Simulations predict *E. coli* expresses *acrAB* in heterogeneous pulses in the absence of stress. Qualitative plots displaying *acrAB* “on”/“off” (dark/light grey) dynamics for 20 randomly selected individual *E. coli* simulations, when exposed to a (i)-(iii) short or (iv)-(vi) long stressor period. Panels display *acrAB* dynamics between timesteps *t* = 120-250 at low, intermediate, and high energy availability. The short stressor period is a single timestep pulse activated at *t* = 150 and the long stressor period is activated at *t* = 150 until *t* = 174.

### 2.4 Efflux network behaviour with a maximal cellular energy budget after exposure to a stressor

We next asked how exposure to a stressor affects network behaviour. Following a simulated stress-free period as in the previous section, we simulated exposure to a stressor for either a short (one timestep) or long (25 timesteps) stress period. In response to stress, the mean expression levels of all genes in the system change, with different rates and magnitude (Fig. 2). *acrAB* mean expression rises throughout stress exposure, thus becoming higher for the longer stress duration (Fig. 2 (i) and (iii)). After the long stress period, the high energy population displays homogeneous behaviour (active efflux expression), increasing the capacity of each cell to remove the stressor (see Fig. 3 (iv) to qualitatively observe this behaviour). The high energy population can then rapidly, and collectively, expel the noxious substance, promptly restoring baseline conditions in mean expression following the steps discussed previously.

When stimulated by a short stress (Fig. 2 (i)), high variability is observed in efflux pump activity throughout the entire stress response, corresponding to diverse responses across the bacterial population. Individual simulation dynamics (Fig. 3 (i)) suggest this behaviour originates from sub-populations of cells developing unsynchronised responses to stress, and a delayed development of the efflux response. The unsynchronised dynamics prolong the population-level stress response, potentially benefiting a clonal population by providing a hedging mechanism against future stress.

Once the simulated stressor is removed, repressive transcription factors resume down-regulation of their GRN target, initially reducing the mean expression level of *marRA* and *acrR*, followed by a reduction in *acrAB* mean expression (Fig. 2). This process switches off the response cascade, allowing *acrAB* mean expression to return to the pre-stress population equilibrium (Fig. 2).

We next asked when the population of *E. coli* cells reached an equilibrium in *acrAB* expression after each perturbation – that is, when the statistics of efflux gene expression levels no longer changed with time in the population after the stress period. We found that the number of timesteps for *acrAB* to attain a post-stress population equilibrium was similar after a short or long stress (9 timesteps vs 8 timesteps respectively, Fig. 2 (i) and (iii)), and the equilibrium eventually attained was the same for both stress durations.

To understand these dynamics in more depth, we next asked how the different patterns of gene expression between which a network can transition vary when a cell is exposed to stress (reflecting the diversity of behaviours supported by the network). We define ‘accessible transitions’ to be the set of expression patterns that can be reached directly from a given network state. To explore this, we analysed the transition matrices describing the dynamics of the *E. coli* regulatory network in the presence and absence of a stressor, visualised through heatmap images (Fig. 4; see Fig. 1(A) for illustration of the process to yield these outputs).

**Figure 4:**
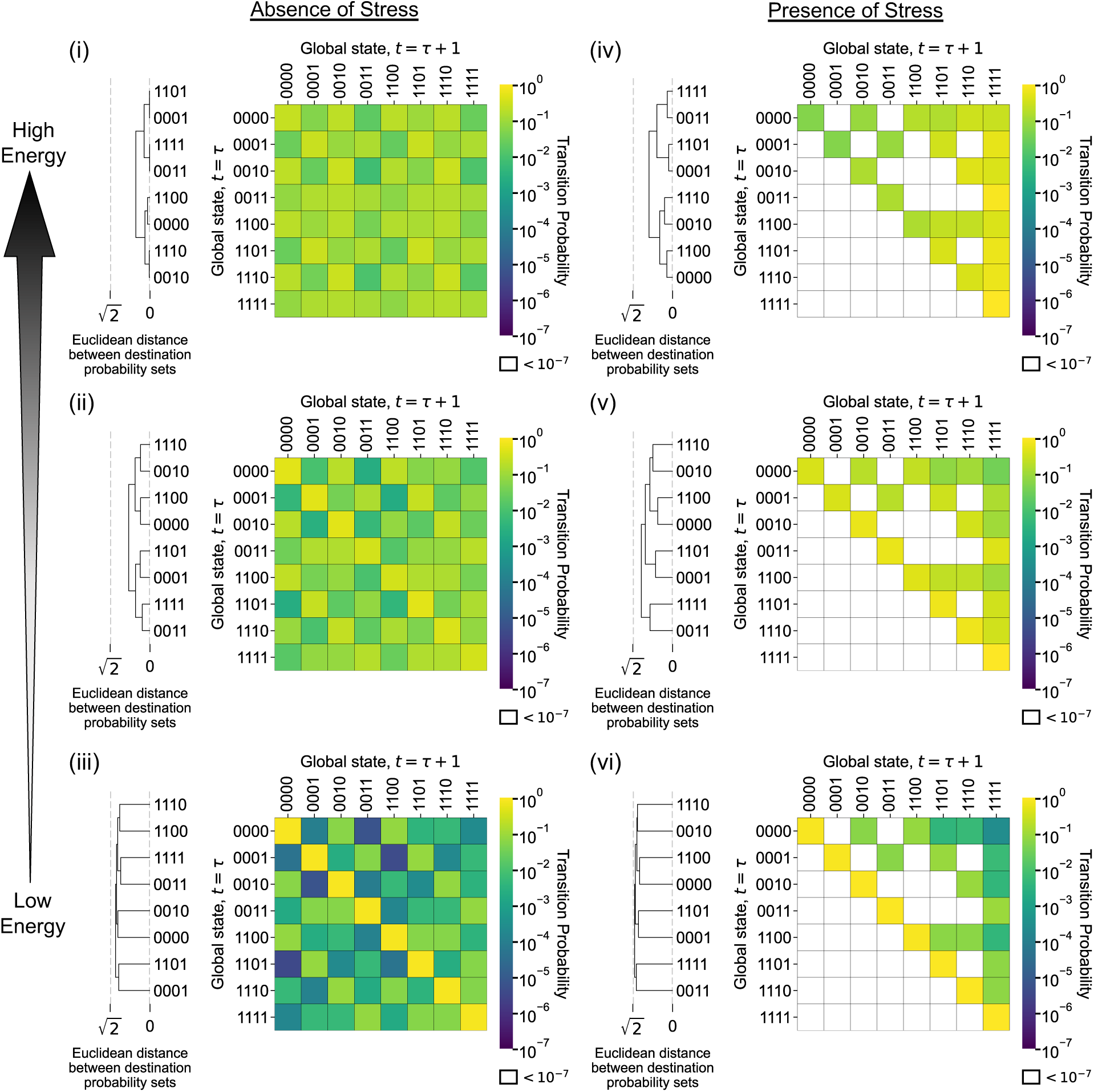
*E. coli* transition dynamics depend on cellular energy and the presence of stress. Heatmap plots showing *E. coli* ’s transition matrix. Each row contains the transition probability distribution from each of possible global states (rows) to each subsequent state (columns), in the absence (i)-(iii) and presence (iv)-(vi) of a stressor, with increasing energy availability; here, due to the coupling of *marR* and *marA* (as they form the *marRA* operon), we do not artificially implement unattainable network states, for example, ‘1011’. Each matrix element is coloured to indicate the probability *p* (using a logarithmic scale) that global state A transitions to global state B from timestep *t* = *τ* to *t* = *τ* +1. Hierarchical clustering of global states is calculated using the destination possibilities (*t* = *τ* +1) for each *t* = *τ* global state, with clustering employing a Euclidean distance metric to determine the distances between each element. Global state binary vectors are displayed in the order [*marR*, *marA*, *acrR*, *acrAB*].

In the absence of stress, all transitions are accessible (Fig. 4 (i)). When a stress is applied, the accessible transitions for each network state are significantly restricted (compare Fig. 4 (i) to (iv)). The presence of a stressor removes the negative regulation within *E. coli* ’s regulatory architecture (Fig. 1(B)), and consequently the ability to actively switch components “off”. The stressed network contains a steady-state attractor – a single state that when reached stays constant – corresponding to all components of the network being “on”. Under stress-free conditions no such single steady-state attractor exists (Fig. 4 (i)). The presence of this attractor implies that an *E. coli* cell under stress shifts its regulatory poise to allow the expression of each GRN component, allowing a response to the stressor and a faster dynamic response when the stress is removed.

### 2.5 Efflux network behaviour is modulated by available cellular energy

We next sought to understand how energy availability changes network behaviour in *E. coli* cells, in an environment with and without a stressor. We first explored how the amount of energy available to a cell affects the accessible network transitions. Both with and without stress, the accessible transitions for each network state are the same across all energy levels, but the probabilities of these transitions vary dramatically with energy (Fig. 4). At lower energies, it is more likely that the system remains in the same state from one timestep to the next; increasing energy distributes the probability of each accessible transition more evenly amongst the total accessible transitions.

When a stressor is present, accessible transitions are restricted to subsets of the stress-free case at each energy level (Fig. 4 (i)-(iii) *versus* (iv)-(vi)). As energy increases, the likelihood of transitioning between supported network behaviours expands, transitioning more directly toward the steady-state attractor, ‘1111’. Therefore, increasing energy availability supports more rapid shifting of behaviour in the *E. coli* efflux network (Fig. 4), particularly towards a high efflux state.

Next, we asked whether the equilibrium mean expression of efflux pump genes changed at different energy levels, both in the presence and absence of stress. Perhaps counterintuitively, we found that increasing energy availability decreased the mean expression equilibrium of *acrAB* in simulated stress-free conditions (Fig. 2). In the presence of a stressor, the population response is more rapid and of higher magnitude as energy availability increases (Fig. 2). Simulated single-cell dynamics predict cell-to-cell variability in the time taken to transition to an active efflux expression state during stress exposure in each energy state (Fig. 3). The delayed response is more pronounced as energy decreases (Fig. 3 (iv) *versus* (vi)). Following either stress period, the network returns to the pre-stress equilibrium more rapidly at higher energy levels (Fig. 2).

### 2.6 Energy levels influence heterogeneity in individual expression level dynamics

To investigate how energy level influences the detailed dynamics of single-cell pulses of efflux pump genes *acrAB*, we calculated the mean pulse length in *acrAB* expression at different energy levels (Fig. 5). Intuitively, from time-scaling effects, mean pulse length increased as energy decreased (Fig. 5 (i)), agreeing with qualitative observations (Fig. 3 (i)-(iii)). The introduction of a short stress results in only a very small transient increase in mean pulse length, more pronounced at higher energy (Fig. 5). Pulse lengths during the long stress period are more dependent on energy level, with pulse statistics in high-energy cells changing more dramatically than in low-energy cells.

**Figure 5:**
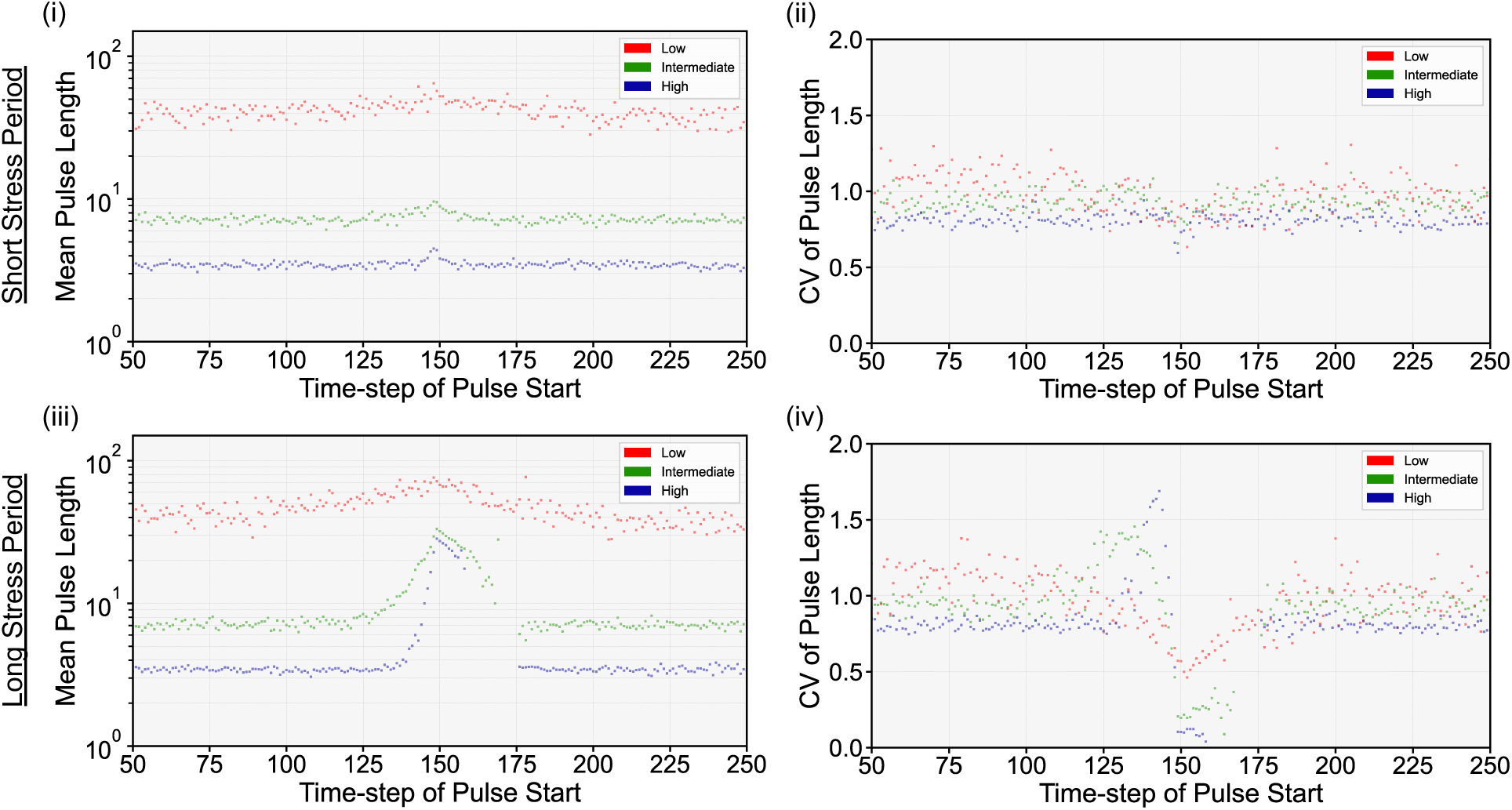
Energy availability and stress modulate *E. coli acrAB* pulse variability. Scatter plots displaying the (logarithmic) mean and coefficient of variation of *acrAB* pulse lengths (coloured dots for low (red), intermediate (green) and high (blue) energy levels) starting at each timestep in *E. coli*. Statistics are displayed between timesteps *t* = 50-250 to observe pulse behaviour before, during and after a (i)-(ii) short or (iii)-(iv) long stress period. The short stressor period is a single timestep pulse activated at *t* = 150 and the long stressor period is activated at *t* = 150 until *t* = 174.

We next addressed our central hypothesis, whether energy availability could be a cause of the observed cell-to-cell variability in efflux pump expression. To explore this, we calculated the dynamics of the expression level coefficient of variation (ELCV) for *E. coli acrAB* at each energy level. As seen previously, substantial cell-to-cell variability exists in the system in the absence of a stressor, reflecting asynchrony and intrinsic noise in the system (Fig. 2). This variability is higher for low-energy cells (ELCV 1.34 at low energy, 1.01 at high energy). Exposure to stress reduces this variability, as the population synchronises and responds in a similar way to the imposed stressor. Exposure to a longer stress period imposed a more substantial decrease in efflux variability than a shorter stress at all energy levels (Fig. 2 (i) and (iii)) and, as observed for the maximal energy case (Section 2.4), the ELCV converged to zero at intermediate to high energy (Fig. 2 (iii)). After the stress exposure, the original level of variability is recovered, allowing the same potential ‘hedging’ against further stress in the future (Fig. 2 (i)) [55–58].

Next, we quantitatively investigated how energy level influences the variability of efflux pump gene expression pulses, by calculating the coefficient of variation of pulse lengths (PLCV) starting from each timestep in *acrAB* (Fig. 5). In the absence of stress, heterogeneity in pulse length is higher at lower energy (Fig. 5 (ii) and (iv)), whilst high energy levels display relatively low PLCV *<* 1, corresponding to lower variability when dynamics are faster and individual events less rare. As with expression level dynamics, when stimulated by a short stress period, PLCV decreases, more dramatically at lower energy levels (Fig. 5 (ii)), as the population becomes more synchronised by the imposition of an external stress. For longer stress periods (Fig. 5 (iv)), differences between different energy levels are more dramatic. Pulse lengths remain highly variable for the low energy population, but PLCV decreases to zero at higher energy. Thus, higher energy availability allows more concerted and synchronous population-level behaviour in dealing with the stressor. Following stress exposure, PLCV behaviour returns to its pre-stress level for each energy level.

### 2.7 *E. coli* and *Salmonella* efflux networks share functional similarity

We next compared the behaviour of the *Salmonella* network to identify similarities and differences between bacterial species. We first explored the capacity of the *Salmonella* network to transition between expression states. As with *E. coli*, the *Salmonella* network architecture supports the same accessible transitions (Supplementary Fig. S5). Clustering by the set of accessible transitions for each global state at a given energy level shows analogous features in both bacterial species, with transitions between states limited by energy (Supplementary Fig. S5).

Next, we asked how *Salmonella*’s *acrAB* efflux pump genes respond to a stressor, compared to the homologous efflux pump in *E. coli*, by analysing the population-level activity of *acrAB* in both species in response to a short and long stress period (Fig. 2 and Supplementary Fig. S3 for *E. coli* and *Salmonella* respectively). Simulation results predict both species respond more dynamically to stress at higher energy, regardless of stress duration and, at each energy level, rapidly return to baseline conditions after the stress period (compare Fig. 2 and Supplementary Fig. S3).

*Salmonella* displays several other analogous behaviours compared to *E. coli*. At maximal energy availability, the number of timesteps for *Salmonella* to return to the mean expression baseline after a short and long stress period is similar (Supplementary Fig. S3), as observed for *E. coli* (see Section 2.3). Other analogous behaviours include a zero *acrAB* ELCV at the maximum stress response to the long stress period (Supplementary Fig. S3 (iii)), and the observation of heterogeneous pulses in network components (Supplementary Figs. S7-S8).

Simulations predict the stress-free *acrAB* ELCV is lower in *Salmonella* at low and intermediate energy, but greater at high energy, compared to *E. coli* (compare Fig. 2 (i) and Supplementary Fig. S3 (i)). This behaviour is a downstream consequence of the mean and variability in global activator expression in each species, which is modulated by the energy level, and the decoupled expression of *Salmonella*’s activator *ramA* and repressor *ramR*, compared with *E. coli* ’s coupled *marRA* operon (Fig. 1(B)-(C)).

Together, this suggests the gene regulatory networks for both bacterial species in this study display behavioural similarities in the presence and absence of a stressor, but despite structural similarities in their architectures (Fig. 1) there are differences in the dynamics of efflux pump expression (Fig. 2 and Supplementary Fig. S3).

## 3 Discussion

We have applied a Boolean modelling framework to coarse-grained *E. coli* and *Salmonella* efflux pump gene regulatory networks, to explore how efflux pump activity depends on the energy available to fuel the expression of efflux proteins and their regulators. We find quantitative support for our hypothesis that differences in available energy can contribute to cell heterogeneity in efflux pump expression, and dissect this heterogeneity in different mechanisms (see Table 1 for a summary).

**Table 1:**
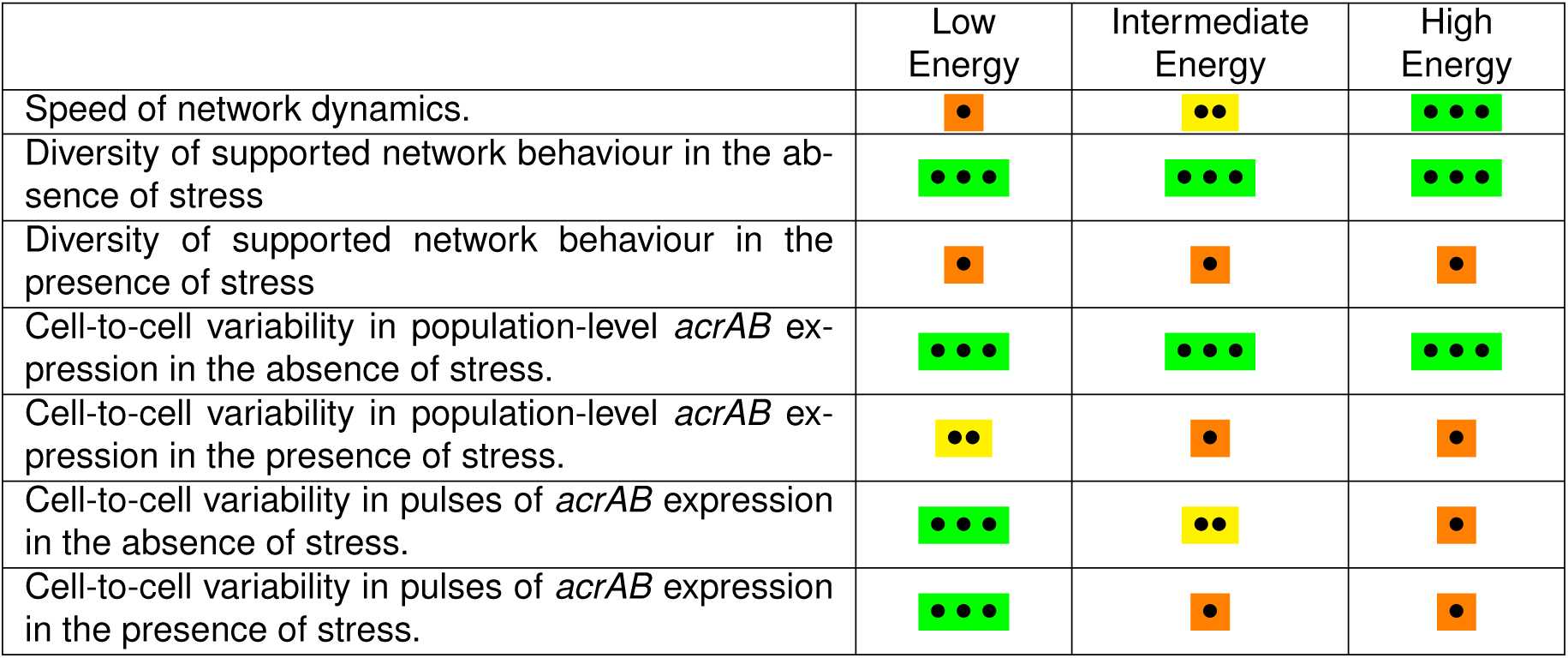
Summary of energy availability effects on regulatory network behaviour. The table displays a summary of our main results from applying the modified Boolean network framework to E. coli and Salmonella acrAB GRNs. The effect of low, intermediate, and high energy availability on each key result is presented through a discretised scale: low (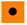), medium (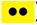) and high (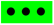).

|  | Low Energy | Intermediate Energy | High Energy |
| --- | --- | --- | --- |
| Speed of network dynamics. | ● | ●● | ●●● |
| Diversity of supported network behaviour in the absence of stress | ●●● | ●●● | ●●● |
| Diversity of supported network behaviour in the presence of stress | ● | ● | ● |
| Cell-to-cell variability in population-level *acrAB* expression in the absence of stress. | ●●● | ●●● | ●●● |
| Cell-to-cell variability in population-level *acrAB* expression in the presence of stress. | ●● | ● | ● |
| Cell-to-cell variability in pulses of *acrAB* expression in the absence of stress. | ●●● | ●● | ● |
| Cell-to-cell variability in pulses of *acrAB* expression in the presence of stress. | ●●● | ● | ● |

**Table 2:** E. coli node initial values. Table defines the stressed and unstressed initial conditions for E. coli gene regulatory network components (Fig. 1(B)).

|  | *marR* | *marA* | *acrR* | *acrAB* |
| --- | --- | --- | --- | --- |
| Unstressed | 0 | 0 | 1 | 0 |
| Stressed | 1 | 1 | 0 | 1 |

**Table 3:** *Salmonella* node initial values. Table defines the stressed and unstressed initial conditions for *Salmonella* gene regulatory network components (Fig. 1(C)).

|  | *ramR* | *ramA* | *acrR* | *acrAB* |
| --- | --- | --- | --- | --- |
| Unstressed | 1 | 0 | 1 | 0 |
| Stressed | 0 | 1 | 0 | 1 |

In the absence of cell-to-cell energy variability, our model suggests (supported by experiments) that substantial intrinsic variability exists in efflux gene expression. The imposition of a population-wide stressor reduces this intrinsic variability by partially synchronising cell behaviour; stochastic effects after stress exposure recover the original variability. This intrinsic noise can potentially be acting to hedge the population against future stress, while maintaining cells that invest less resource in defence mechanisms and/or protect the lineage from diverse environmental conditions. When extrinsic differences in ATP availability between cells are additionally imposed, the population variability in efflux response will be amplified. Cells with lower energy levels will experience more variable expression of efflux components (amplifying intrinsic variability), slower responses to stressors, and hence slower removal of stressors from the environment. In future, experimental tests of this hypothesis could involve, for example, jointly tracking bacterial ATP levels and efflux gene expression in a culture prior to, during and post antibiotic stress using fluorescent reporters [11, 12, 16, 59].

In our model, stress-free bacteria naturally express heterogeneous pulses of efflux genes (Fig. 3 and Supplementary Fig. S7). This behaviour may be biologically valuable for several reasons. The products of *acrAB* have an extremely long half-life in *E. coli* [60], suggesting that the genes may only need to be transcribed in short pulses (which is predicted to be a less energetically demanding process compared to translation [61]) for the protein to be in an abundant supply. Dealing with external stressors is one role of the AcrAB-TolC efflux pump, but it also, for example, removes endogenous waste products [62], suggesting the pulse behaviour could aid physiological functions by expressing *acrAB* mRNA to be used in these processes. The pulse behaviour may also confer a natural resistance phenotype to a fraction of cells in stress-free environmental conditions at all times, as reported experimentally in *Salmonella enterica* [13]. Therefore, the predicted pulsing behaviour may contribute to several essential intracellular functions.

Recent mathematical models have studied the *mar* network at the protein level, suggesting the motif is monostable, and supports a single deterministic steady-state in the absence of stress [63, 64]. A theoretical study has also shown, at the protein level, the development of stochastic pulses in *marRA* products [64]. The cell-to-cell heterogeneity in *mar* expression that we predict agrees with ODE models of the *mar* operon – which account for the inherent stochasticity of biological systems – and experimental demonstrations of heterogeneous cell-to-cell steady-state *mar* expression within a population, in the presence and absence of stress [64–66] (Fig. 2 and Supplementary Fig. S6). With the development of advanced experimental assays, future studies may resolve the behaviour at the transcript level to test our theoretical predictions.

Even for these relatively simple networks, our Boolean modelling approach has demonstrated general principles of energy influence without requiring characterisation of a large parameter space. Developed with a view to scalability, this approach can be applied to many naturally-occurring decision-making phenomena. Our previous work [14] has demonstrated the profound influence that energy variability can have on simple decision-making circuits; this GRN approach allows the expansion of this philosophy to a much wider range of architectures, including the broader regulatory context of these core networks (see Supplementary Information sections 5.1-5.2). As cell-to-cell variability in ATP is increasingly elucidated [18–20] we believe that our approach will help reveal the theoretical underpinnings of this important influence on regulatory dynamics.

Our hypothesised relationship between energy availability and efflux-pump activity, variability, and capacity to respond to antimicrobials has, as far as we are aware, not been previously considered. However, our hypothesis is not restricted to this AMR mechanism or motif. A connected phenomenon that may be mechanistically related is the formation of bacterial persisters in the presence of antibiotics [55, 67, 68]. Here, phenotypic variants display a transient tolerance to external stress [69, 70], causing a leading factor in the recalcitrance of chronic infections [71–74]. An approach grounded in optimal control theory has found that the persister strategy is dependent upon stochastic fluctuations in both the proportion of persisters in the colony and the environment [75]. The results also predict that producing persisters in the absence of environmental volatility leads to a lower per-capita growth rate, giving the population a possible evolutionary disadvantage. Increasing evidence has reported a mechanism linking persister cells and intracellular energy, or more specifically ATP [17, 76–79]. One study has gone further, suggesting there may be a general low energy mechanism of persister formation in bacteria [80]. This relationship bears substantial similarities to our model where we predict high variability and noise in gene expression pulses in low energy cells, compared to intermediate and high energy cells (Fig. 2 and Fig. 5). High expression variability contributes to cell-to-cell variation within a clonal population, implying a different set of phenotypes may be expressed in the low energy sub-population. As energy, or more precisely ATP, increases, cell-to-cell variability decreases, implying a uniform phenotype displayed across the population. We would predict, in agreement with growing evidence, intracellular ATP is a factor in the mechanisms of persister generation, and future work will explore this using our approach when the governing GRNs can be identified.

## 4 Methods

### 4.1 Wiring Diagrams

In the supplement (Supplementary Fig. S1) we present the wiring diagram form of each gene regulatory network (Fig. 1(B)-(C)). The wiring diagrams contain the regulatory interactions in each GRN (activatory or inhibitory) and edge weights, outputting a directed graph. A stress, such as antimicrobial exposure, is included through the ‘stress node’, *S*, within the wiring diagram. A stress input is modelled as repressing gene-gene regulatory edges, representing a ‘repressive’ modulation of transcription factors (MarR/RamR and AcrR). The capacity to expel substances by the efflux pump is integrated through a repressive regulation from *acrAB* to *S*.

Estimates of edge weights were gleaned from the relevant literature, integrating the natural behaviour of each network, and giving each regulatory actor the ability to operate. In *E. coli*, the *marRA* operon is separated into its individual components to allow each regulation of *marRA* to be captured, but are coupled in all iterative updates; the inability to model a node that both positively and negatively regulates itself is a limitation of the Boolean modelling framework. Experimental data has shown transcription of *marA* and *acrA* increase in *marR* mutant cells [81], and MICs of various antibiotics increase in an *acrR* mutant strain [44] (consistent with the limited repressor function of AcrR [41]). Together, these suggest, under under normal conditions, edge weights of interactions from *marR* to *marRA* and *acrR* to *acrAB* should be dominant. Inactivation of the *ramR* gene upstream of *ramA* resulted in increased expression of *ramA* and the AcrAB efflux pump in *Salmonella* [82]. This requires the edge weight from *ramR* to *ramA* to outweigh *ramA* basal expression under normal conditions. Constitutive gene expression is captured using ‘ghost nodes’. These nodes are permanently “on” in the network, supplying a constant positive influx to a node.

Interaction matrix J is constructed from the data contained within a wiring diagram. A weighted edge from node *j* to node *i*, is represented by element J*_i,j_* within J. Each matrix element can take a positive, negative or zero value; edge weight sign corresponds to a negative, positive or absence of regulation respectively. Since edges are directed, J*_i,j_* ≠ J*_j,i_*, except for *i* = *j* or J*_i,j_* = J*_j,i_* = 0. Matrix J is implemented in numerical simulations of the Boolean model and has dimension *N* -by-*N* (*N* is the total number of nodes in a wiring diagram, including ghost and stress nodes).

### 4.2 Initial conditions

Two configurations of initial node states were considered in simulations of the *E. coli* and *Salmonella* wiring diagrams. These were named the “unstressed” and “stressed” initial condition, with the initial node states set as follows:

### 4.3 Modulation of transition rates allows energy variability to be captured in GRN models

Wiring diagrams, consisting of nodes and edges, schematically represent the interactions between elements within regulatory networks (as shown in Supplementary Fig. S1). The edges often illustrate combined processes such as transcription, translation and phosphorylation. However, the information contained within edges in regulatory networks is coarse-grained, omitting a substantial amount of important biological detail. Several of the processes represented by these edges require ATP as an energy source [29, 53], so there exists a core energy dependence in the dynamics modelled within each wiring diagram.

To address this and capture the fact that all the processes involved in gene expression depend on ATP, we constructed a modelling approach modulating the strength of regulation from node *X* to node *Y* according to cellular energy levels. This reflects the fact that, for example, cells with low ATP will have a lower rate of transcription elongation per timestep [53]. We interpret this modulation as influencing the rate with which regulatory interactions can be applied from node *X* to node *Y* . The rationale for this picture is that if the expression of regulatory components is slowed by reduced energy levels, the corresponding biochemical interactions will occur at a lower rate.

In the absence of energy variability, our asynchronous Boolean modelling framework interprets regulatory interactions as stochastic events occurring with a characteristic mean rate of one per timestep per interaction. To include the influence of energy availability on the system, we allow this rate to be a function of energy level. To capture an order-of-magnitude range across cells, we use rates for simulation timestep of 1 for ‘high’ energy (hence, all interactions are always realised), 0.5 for ‘intermediate’ energy, and 0.1 for ‘low’ energy. We thus modulate the probability of any given interaction being realised in a given simulation timestep by the level of available cellular energy.

### 4.4 Node update function

Nodes within the Boolean network are updated iteratively, using an update function. An update function defines the value of each node according to its regulatory nodes and remains fixed for all timesteps [83]. We consider a threshold update function to update the state of node *i* at timestep *t* = *τ* + 1, denoted *σ_i_*(*τ* + 1). The argument of the update function consists of the regulatory nodes of node *i*, the states of these nodes at *t* = *τ* and the *N* -by-*N* interaction matrix J, where *N* is the total number of nodes in the wiring diagram. The summation value determines the state of node *i* at *t* = *τ* + 1.

During an active stress time frame, the state of stress node *S* (Supplementary Fig. S1) is imposed to be “on” at the start of each timestep, regardless of the behaviour from the previous update. Once the final timestep within the stress period is reached, *S* is allowed to be switched “off” from regulation within the network.

We consider the update function displayed in Equation (4.1). Here, a node requires a non-zero sum of its inputs to change state from *t* = *τ* to *t* = *τ* + 1. In the case where the sum is zero, the node state remains unchanged.

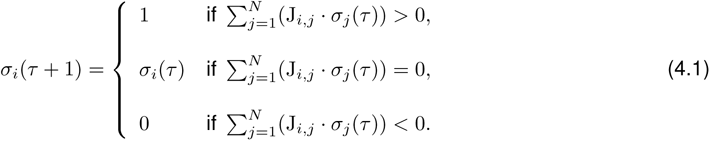

Alternate threshold update functions are possible. We could, for example, consider the conditions that set a node as “on” if there is an overall positive input, and “off” otherwise. However, this would automatically remove a stress in the system by setting the stress node state to “off” once reaching the end-point of the stress time frame. This removes the capability of the efflux node to ‘expel’ the stress through feedback to the stress node *S* (Supplementary Fig. S1), behaviour which is not the focus of this study. By choosing update rule (4.1), it allows the efflux pump node to have the capacity to remove stress in the network, through regulatory interactions, depicting the behaviour observed in bacteria.

At each timestep, the binary vector containing all node states is the global, or network, state [84]. The set of all possible global states forms the state space of a network [83]. The state space for each network in this study is of size 2*^M^* for *Salmonella*, as we are interested in the dynamics of the *M* -component GRN network being modelled, and 2*^M−^*^1^ for *E. coli*, due to the coupling of *marR* and *marA*.

### 4.5 Node update methods

There are two methods for updating global states, synchronous and asynchronous. In the synchronous mode, all the nodes (including the stress node) are updated at every timestep, with the state of each node at *t* = *τ* + 1 depending on the states of its inputs at *t* = *τ* . The deterministic nature of synchronous Boolean models, combined with each model containing a finite state space (2*^M^* or 2*^M−^*^1^), where *M* is the number of elements in the original GRN, guarantees that a synchronously updated time series simulation will eventually end at a steady-state attractor, or a cycle of repeating global states [83]. It also means the network will always reach the same state for a given initial condition and number of timesteps. We do not consider synchronous updating in this study due to the artefacts that can arise from its imposition, and its inability to model stochastic processes. A more complex, but biologically realistic, protocol is the asynchronous mode, which gives a reasonable overview of the random dynamics in the cell and better captures transient behaviour. This method updates nodes of the system according to the last update of their regulator nodes, either from the previous or current iteration [85]. Previous studies have employed numerous different versions of asynchronous updating [27, 86–88], including methods where all nodes are updated according to a random sequence, or one randomly selected node is updated at a time step [89]. In this study the asynchronous protocol updates *η* nodes per timestep, with *η* drawn from a Poisson distribution with a mean equal to the number of original elements in the regulatory network (*M* ), plus the stress node. Additionally, each node can be updated more than once per timestep and the state of each node at *t* = *τ* +1 depends on the states of its inputs at the most recent update. We used an asynchronous protocol because we believe it better captures the stochasticity in the cell.

### 4.6 Boolean Analysis

Taken together, the wiring diagram, initial conditions, update functions and update method form the Boolean network model. Time-dependent numerical solutions of the presented Boolean model were generated in Python 3.8.2 using modules *numpy*, *pandas* and *random*. Numerical solutions were calculated from prescribed initial conditions covering the stressed and unstressed state of each network. Time-series figures, displaying the expression profiles of each node, were constructed from the mean of node expression and standard deviation per timestep, from 10^4^ simulations. Visualisations of ‘single-cell’ expression profiles, and pulse statistics, were constructed from 20 and 5000 randomly selected simulations respectively. Transition matrices were constructed from the simulation data and visualised through a heatmap structure, from 10^7^ simulations. Each entry of the transition matrix is a non-negative real number representing the probability of transitioning between two network states from timestep *t* = *τ* to *t* = *τ* + 1. Clustering of global states was performed using the same data, with a Euclidean distance matrix from modules *scipy.cluster* and *scipy.cluster.hierarchy*. All scripts used in this study are openly accessible through https://github.com/StochasticBiology/boolean-efflux.git.

## Acknowledgements

RK thanks the Wellcome Trust for funding (grant reference 108876/Z/15/Z).

## 5 Supplementary Information

### 5.1 A Bottom-up Construction of Efflux Pump Gene Regulatory Networks

A key regulator of efflux-related drug resistance in *E. coli* is the multiple antibiotic resistance (*mar* ) operon [1, 2]. This genomic region contains the *marRAB* operon, encoding three proteins named MarR, MarA and MarB [1]. MarR has a repressive function, acting on the basal expression of the *mar* operon in stress-free conditions [1, 3, 4]. MarA, the *marA* gene product, is a member of the AraC/XylS family of proteins which induces *acrAB* expression and undergoes positive feedback through auto-activation of the *marRAB* operon [5, 6]; in addition to *acrAB* and *marRAB*, MarA regulates the expression of numerous other genes, either directly or indirectly, in its regulon [7]. The final product from the operon, MarB, is a small periplasmic protein that represses *marRAB* but does so through an unknown mechanism and is therefore omitted from our regulatory network [8].

*Salmonella* has a second MarA-like protein called RamA, an AraC/XylS transcription activator synthesised from *ramA* [9]. *ramA* is a homologue of the *E. coli marA* gene, having 45% similarity [10]; while not present in *E. coli*, *ramA* is present in a wide range of Enterobacteriaceae. Like MarA in *E. coli*, RamA positively regulates *acrAB* expression in *S. enterica* [11, 12] by binding to the region upstream of *acrAB*. RamR is the transcriptional repressor of *ramR* and *ramA*, binding to the *ramRA* intergenic region [13, 14]. We include the *ramRA* regulatory region in *Salmonella*’s gene regulatory network (Fig. 1(C)) and omit the *mar* operon due to previous studies showing inactivation of *marR* and *marA* (as well as *soxR*, and *soxS*) did not affect the susceptibilities of a *S. enterica* serovar Typhimurium strain, whereas inactivating *ramR* resulted in a multidrug resistance phenotype [13]. This observation, along with other supporting studies, suggests the *ramRA* regulatory region primarily controls AcrAB and efflux-mediated multidrug resistance in *Salmonella* [11, 15]. The major roles of the *mar* operon and *ramRA* region in regulating *acrAB* expression, in *E. coli* and *Salmonella* respectively, supports the choice of these regulators in our coarse-grained GRNs (Fig. 1(B)-(C)). Additional regulation of *acrAB* is provided locally in both species by *acrR* [16, 17]. Its product, AcrR, is known to repress both its own synthesis and *acrAB* from a divergently transcribed region [16]. Under normal conditions, *acrR* is transcribed at a basal level, preventing over-expression of *acrAB* [16, 18]. The expression of efflux pumps is inducible by the presence of environmental signals, or chemical compounds (such as antibiotics). The inducibility of efflux genes enables bacteria to adapt to a wide range of environmental conditions. Noxious substances or stressors can bind to transcription factors MarR, RamR and AcrR. In *E. coli*, derepression of the *mar* operon can occur through stress molecules reversibly binding to MarR, inhibiting its ability to bind to DNA and interfering with the repressor activity of MarR [19, 20]. Derepression of *mar*, by the MarR inhibitor salicylate for example [21], promotes increased transcription of *marRAB*, in particular *marA*, and consequently *acrAB* [19, 20]. The activity of AcrR, which belongs to the TetR family of transcriptional repressors, has been observed to be induced by ligands such as rhodamine 6G, ethidium, and proflavin [22]. In *Salmonella*, substances increase *acrAB* expression through the increased expression of both *ramA* and *ramR*, by inhibiting the binding of RamR to the P*_ramA_* region [14, 23]. Studies have shown that not all antibiotics significantly increase the expression of *ramA* [24], suggesting that not all efflux substrates are molecules that induce efflux expression. In our model we consider the effect of molecules that induce efflux expression and can be exported by the efflux pump, displayed through a repressive interaction from *acrAB* to a stress node *S* in Supplementary Fig. S1.

### 5.2 Additional Actors in Gene Regulatory Networks

The regulatory models considered in this study focus on transcript level behaviour, excluding additional regulatory processes, such as Lon protease and CsrA, acting at the post-transcriptional or post-translational level to fine-tune the regulation of *acrAB* expression [25–27]. Additional layers of control from, for example, *soxRS*, *rob* and *acrZ*, are also known to be integrated into regulating AcrAB expression [28–31]. These regulators are thought to play a minor role in *acrAB* regulation, or efflux-related resistance, compared to the *mar*, *ram* and *acrR* regulators [11, 13, 18, 29, 32, 33].

Feedback between efflux genes and transcriptional regulators has also been identified [24, 34–36], however, full knowledge of their mechanism, including whether it is direct or indirect, is still incomplete; in the case where the relationship between the regulatory players is stated, but the potential interactions are still unknown, they are omitted from the GRNs. Further, bacterial efflux pump regulation is highly interlinked, with various efflux pumps previously shown to display compensatory expression when alternative efflux pump genes are up- or down-regulated. In *Salmonella*, for example, expression of *acrD* and *acrF* genes, encoding homologous RND pumps in enterobacteria, were significantly up-regulated in an *acrB* knockout mutant [35–37]. Likewise, analogous behaviour is observed for *acrB* expression when Δ*acrF* mutants, or

Δ*acrB* Δ*acrF* double mutants are considered [35]. AcrAB-TolC is a clinically relevant multidrug resistance efflux pump within *E. coli* and *Salmonella* [38–40]. The key involvement of AcrAB in clinical resistance corroborates the focus of our study on the regulation of this efflux pump component.

### 5.3 Validating the Theoretical Model Through Independent Experimental Observations

Supplementary Table S1 contains multiple independent experimental observations captured by our theoretical model, and used to validate our mathematical approach. weights.

### 5.4 *Salmonella* and *E. coli* wiring diagram schematics

Supplementary Fig. S1 shows the wiring diagram version of *E. coli* (Fig. 1(B)) and *Salmonella* (Fig. 1(C)) efflux pump gene regulatory networks, including the regulatory interactions illustrating inactivation of transcription factors by noxious substances, and the capacity of the assembled efflux pump (AcrAB-TolC) to remove stressors.

### 5.5 Network dynamics from different initial conditions

The first question we addressed with our model was whether the initial state of the network led to different long-term behaviour in *E. coli* and *Salmonella* cells in environments with and without a stress. We define the stressed and unstressed initial conditions (ICs) for our model by considering the active components in bacterial efflux mechanisms in an environment with and without a stressor (see Methods). We then simulated the evolution of each network over a set time period, starting either from an unstressed state or a stressed state applied for 1 or 25 timesteps (modelling different stress conditions). The stressor was applied by perturbing the network after reaching a population equilibrium, and not during the initial transient activity of the network.

We found that the ICs had no influence on the long-term behaviour of either network in simulated stress-free conditions – for example, compare Fig. 2 and Supplementary Fig. S2 before the short stress period in *E. coli* (similarly, compare the pre-stress behaviour in Supplementary Figs. S3 and S4 for *Salmonella*). When a stress is applied to a network, the response of the network and long-term dynamics are unaffected by the ICs. The analogous response to stress for both ICs is not surprising, as each network is perturbed once population equilibrium – which is the same for both ICs – is reached. Therefore, in the absence of energy availability, the choice of IC does not affect the long-term behaviour.

When allowing for energy variability within the model, identical results were observed (compare Fig. 2 to Supplementary Fig. S2 for *E. coli*), supporting the use of the unstressed initial condition within this study.

### 5.6 *Salmonella* network transitions

Supplementary Fig. S5 displays network state transitions for *Salmonella* GRN (Fig. 1(C)) using heatmaps, with and without stress.

### 5.7 Individual Cell Dynamics Display Significant Cell-to-Cell Heterogeneity

Supplementary Figs. S6-S8 display qualitative dynamics of simulated single-cell *marRA* expression in *E. coli*, and *acrAB* and *ramA* expression in *Salmonella*, respectively.

**Figure S1:**
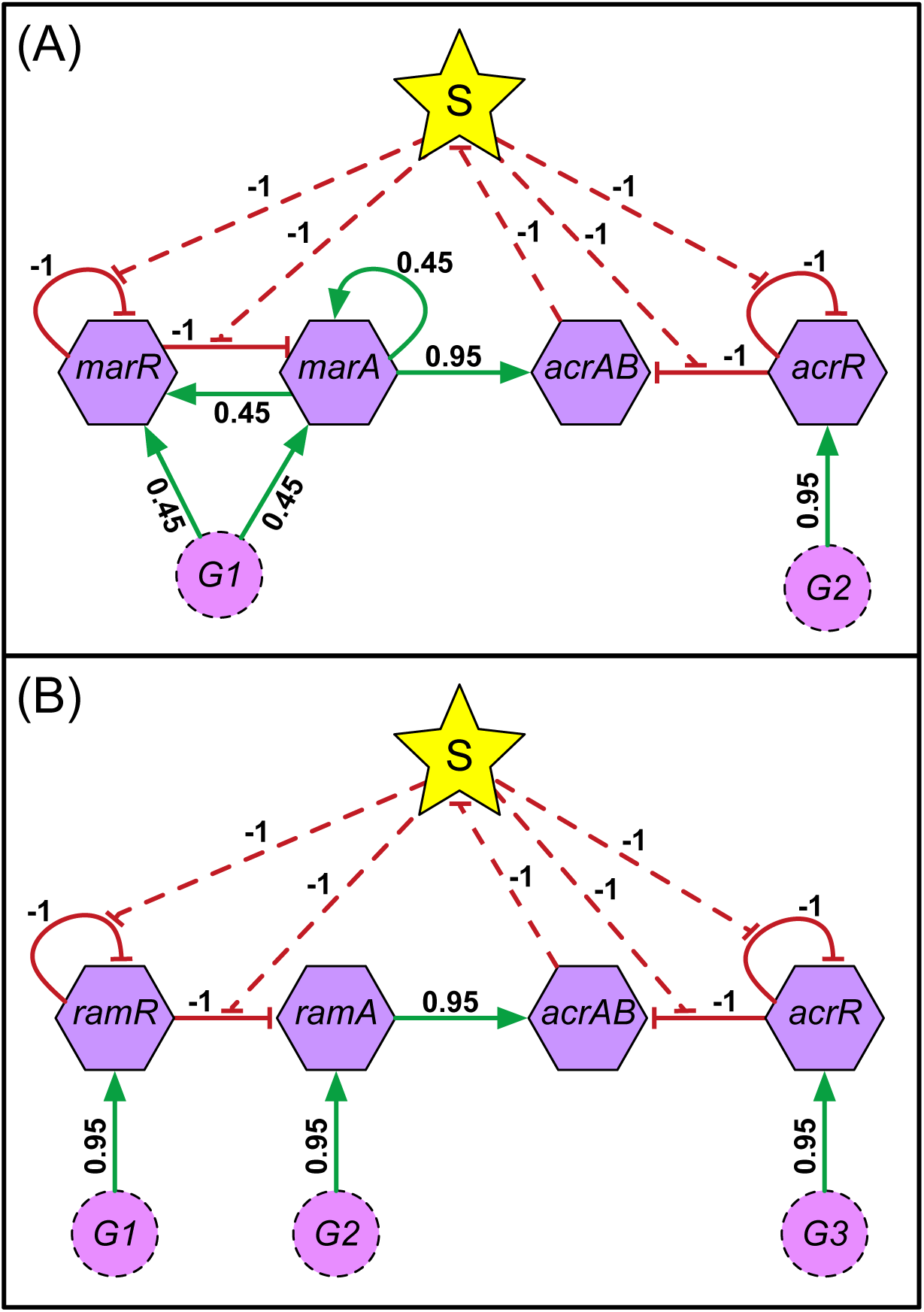
Schematic representation of network wiring diagrams. Wiring diagrams for (A) *E. coli* and (B) *Salmonella* efflux pump GRNs (Fig. 1(B) and (C), respectively). GRN interactions directly modulate network nodes in the wiring diagram depiction, and stress-related regulation (dashed edges to and from node *S*) modulates network regulatory edges; edge weights within the wiring diagram are derived from the literature, and enable each element in the network a chance to act. Each regulatory interaction is colour coded to display positive (green) or negative (red) operations.

**Figure S2:**
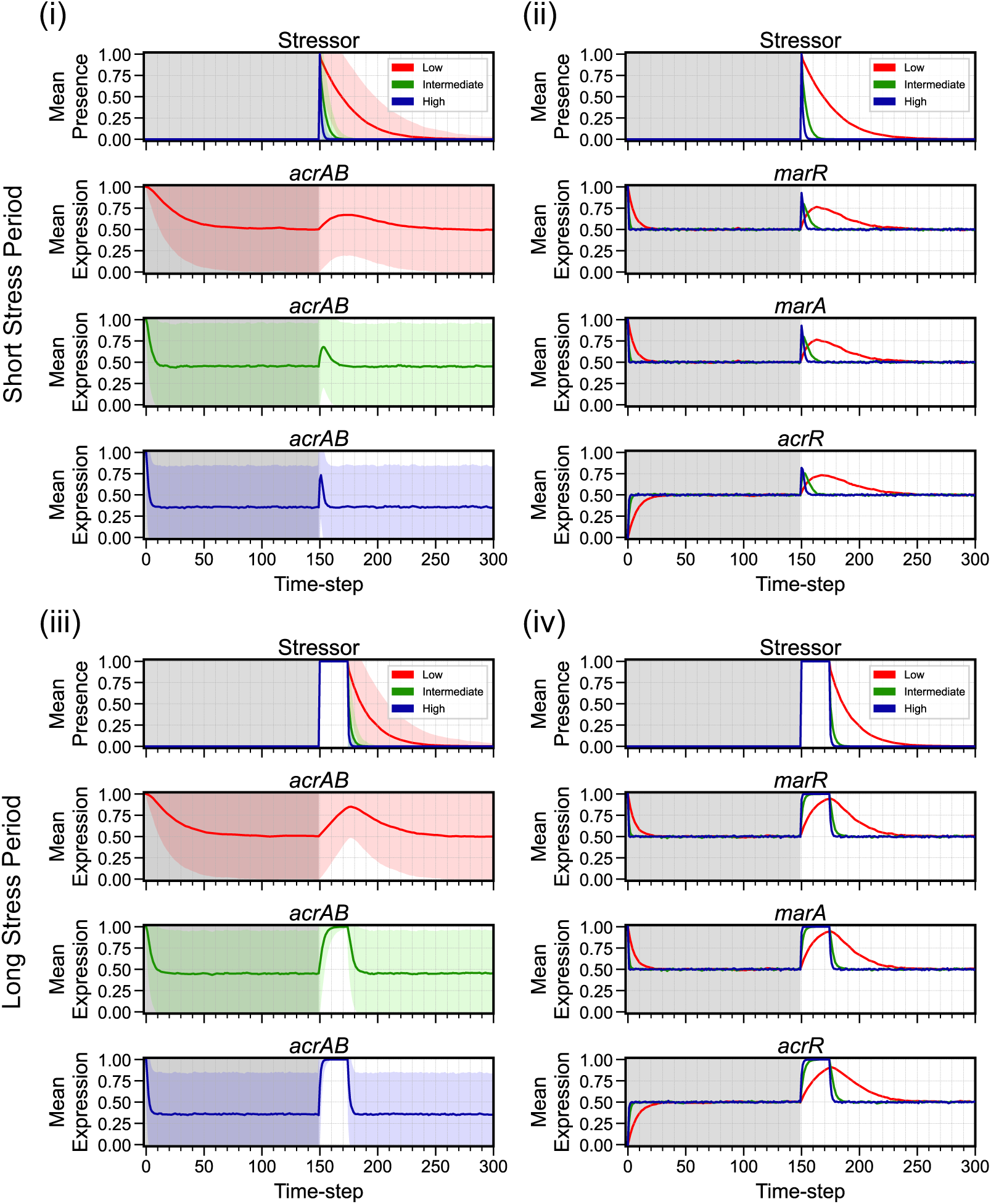
*E. coli* mean expression dynamics are equivalent from different initial conditions. Time-series dynamics of mean expression (coloured line for low (red), intermediate (green) and high (blue) energy levels) for ((i), (iii)) efflux genes *acrAB* and ((ii), (iv)) remaining regulatory components in *E. coli*, from the stressed initial condition; all panels include stressor behaviour. Behaviour is shown when the network is exposed to a (i)-(ii) short (*t* = 150) or (iii)-(iv) long (*t* = 150-174) stress period. The initial time region of each regulatory component prior to stress application (grey shaded region) displays the complete stress-free behaviour. Note: Standard deviation (corresponding coloured shaded area) is included within efflux panels to represent the uncertainty, but the actual expression level cannot exceed the [0, 1] domain.

**Figure S3:**
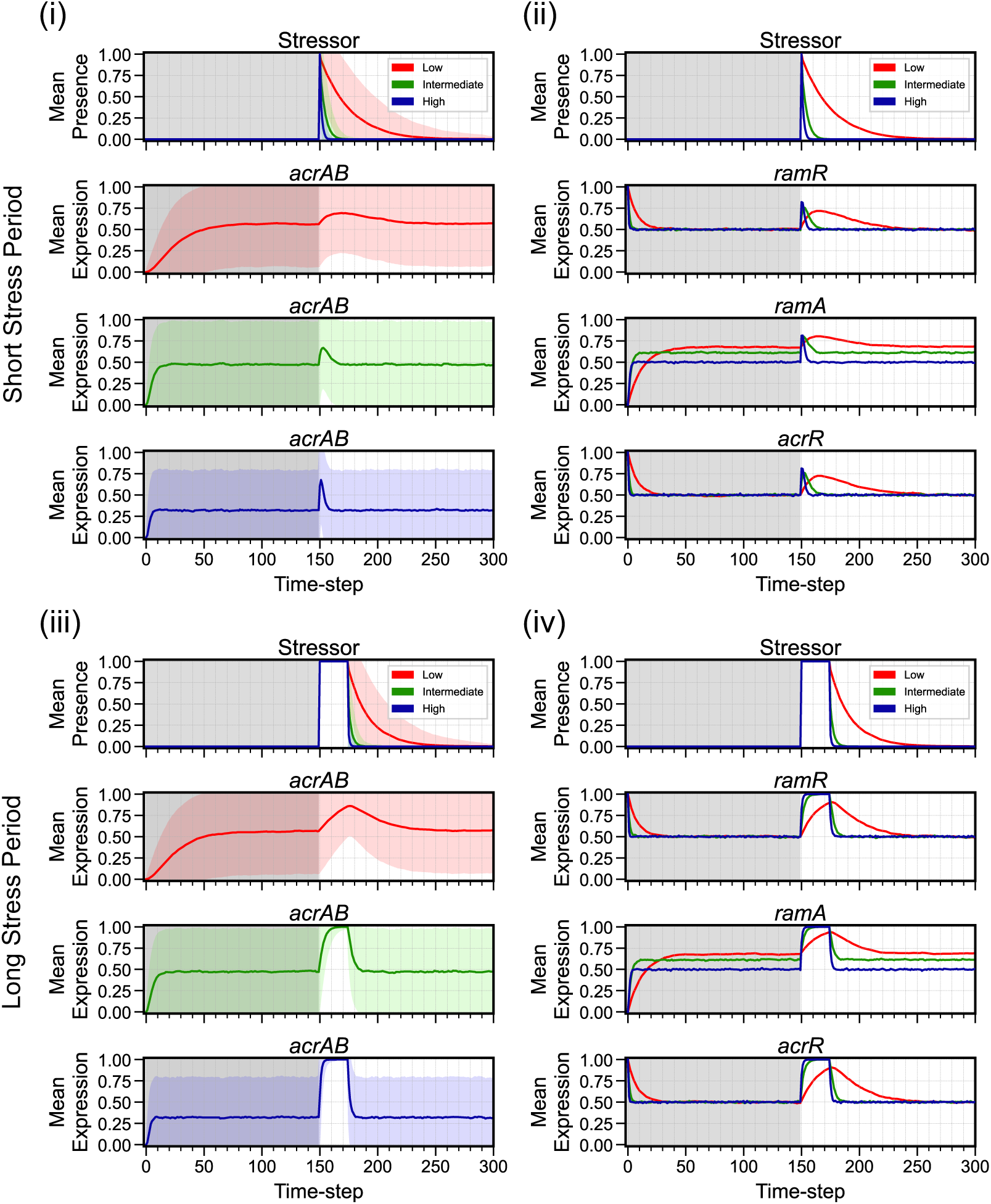
Mean expression dynamics for *Salmonella* from the unstressed initial condition in response to a short or long stressor duration. Time-series dynamics of mean expression (coloured line for low (red), intermediate (green) and high (blue) energy levels) for ((i), (iii)) efflux genes *acrAB* and ((ii), (iv)) remaining regulatory components in *Salmonella*, from the unstressed initial condition; all panels include stressor behaviour. Behaviour is shown when the network is exposed to a (i)-(ii) short (*t* = 150) or (iii)-(iv) long (*t* = 150-174) stress period. The initial time region of each regulatory component prior to stress application (grey shaded region) displays the complete stress-free behaviour. Note: Standard deviation (corresponding coloured shaded area) is included within efflux panels to represent the uncertainty, but the actual expression level cannot exceed the [0, 1] domain.

**Figure S4:**
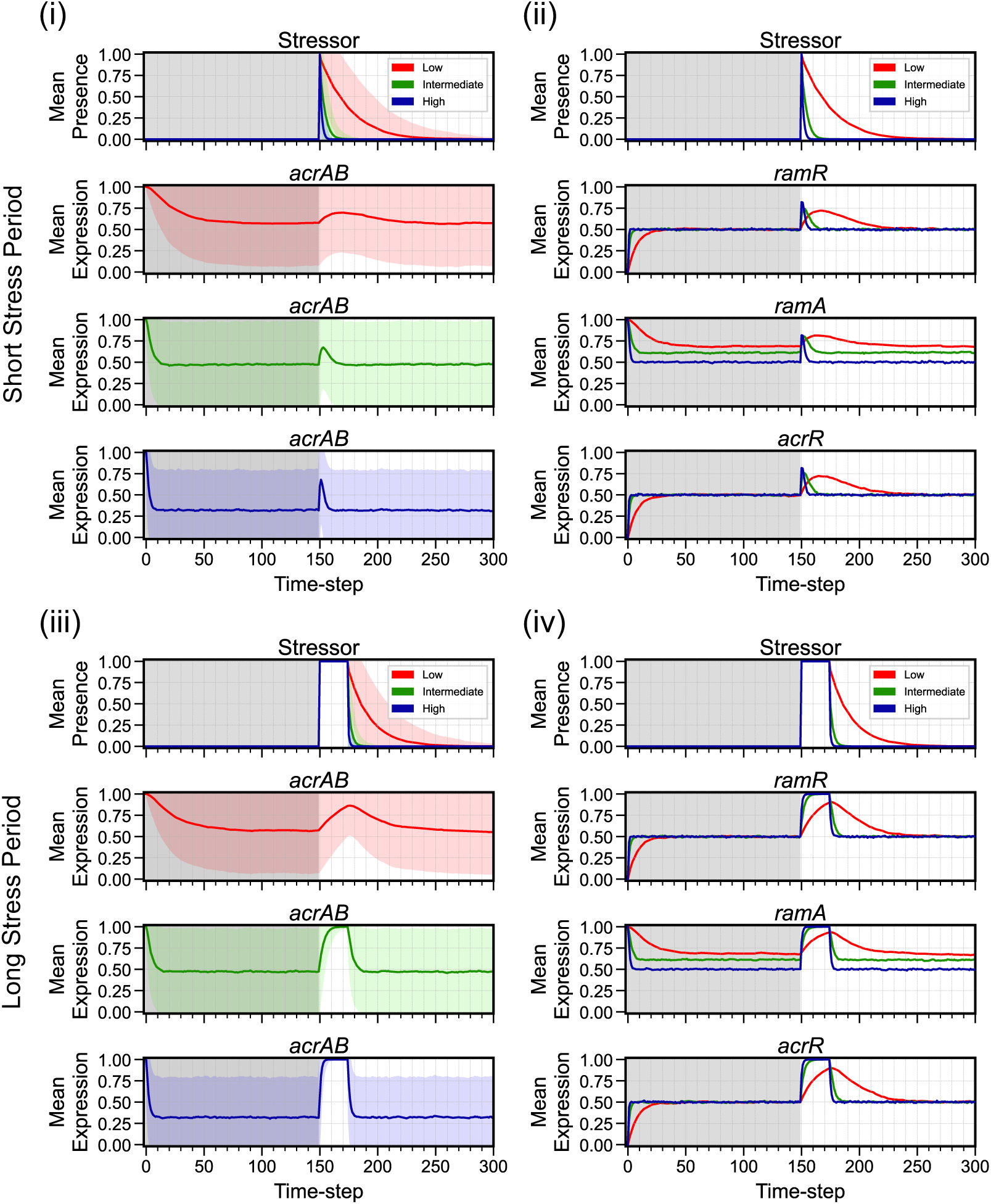
*Salmonella* mean expression dynamics are equivalent from different initial conditions. Time-series dynamics of mean expression (coloured line for low (red), intermediate (green) and high (blue) energy levels) for ((i), (iii)) efflux genes *acrAB* and ((ii), (iv)) remaining regulatory components in *Salmonella*, from the stressed initial condition; all panels include stressor behaviour. Behaviour is shown when the network is exposed to a (i)-(ii) short (*t* = 150) or (iii)-(iv) long (*t* = 150-174) stress period. The initial time region of each regulatory component prior to stress application (grey shaded region) displays the complete stress-free behaviour. Note: Standard deviation (corresponding coloured shaded area) is included within efflux panels to represent the uncertainty, but the actual expression level cannot exceed the [0, 1] domain.

**Figure S5:**
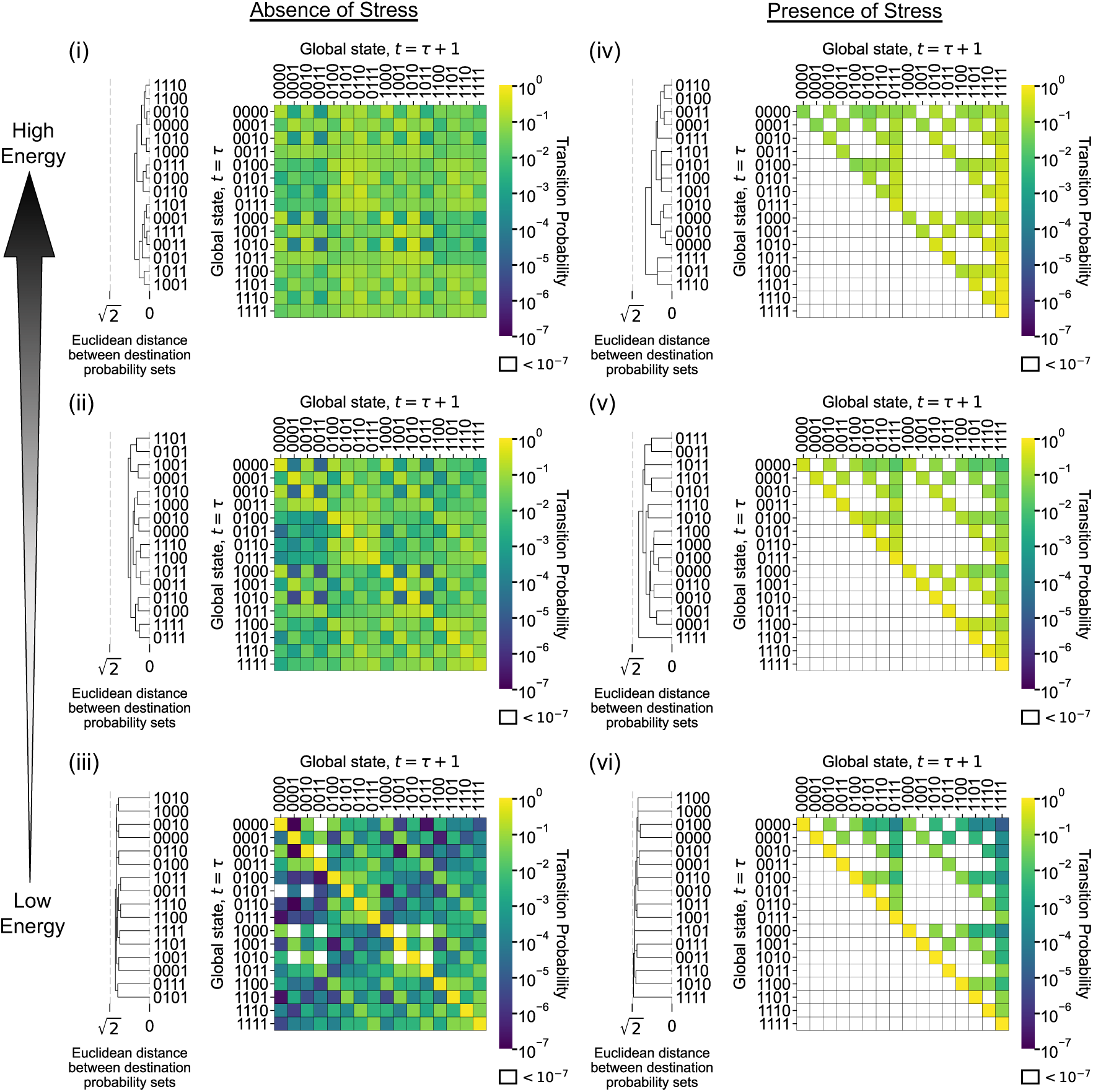
*Salmonella* displays analogous coarse-grained qualitative behaviour compared to *E. coli*. Heatmap plots showing *Salmonella*’s transition matrix. Each row contains the transition probability distribution for each of 2*^M^* possible global states, in the absence (i)-(iii) and presence (iv)-(vi) of a stressor, with increasing energy availability; *M* here is the number of nodes in the initial GRN. Each matrix element is coloured to indicate the probability *p* (using a logarithmic scale) that global state A transitions to global state B from timestep *t* = *τ* to *t* = *τ* + 1. Hierarchical clustering of global states is calculated using the destination possibilities (*t* = *τ* + 1) for each *t* = *τ* global state, with clustering employing a Euclidean distance metric to determine the distances between each element. Global state binary vectors are displayed in the order [*ramR*, *ramA*, *acrR*, *acrAB*].

**Figure S6:**
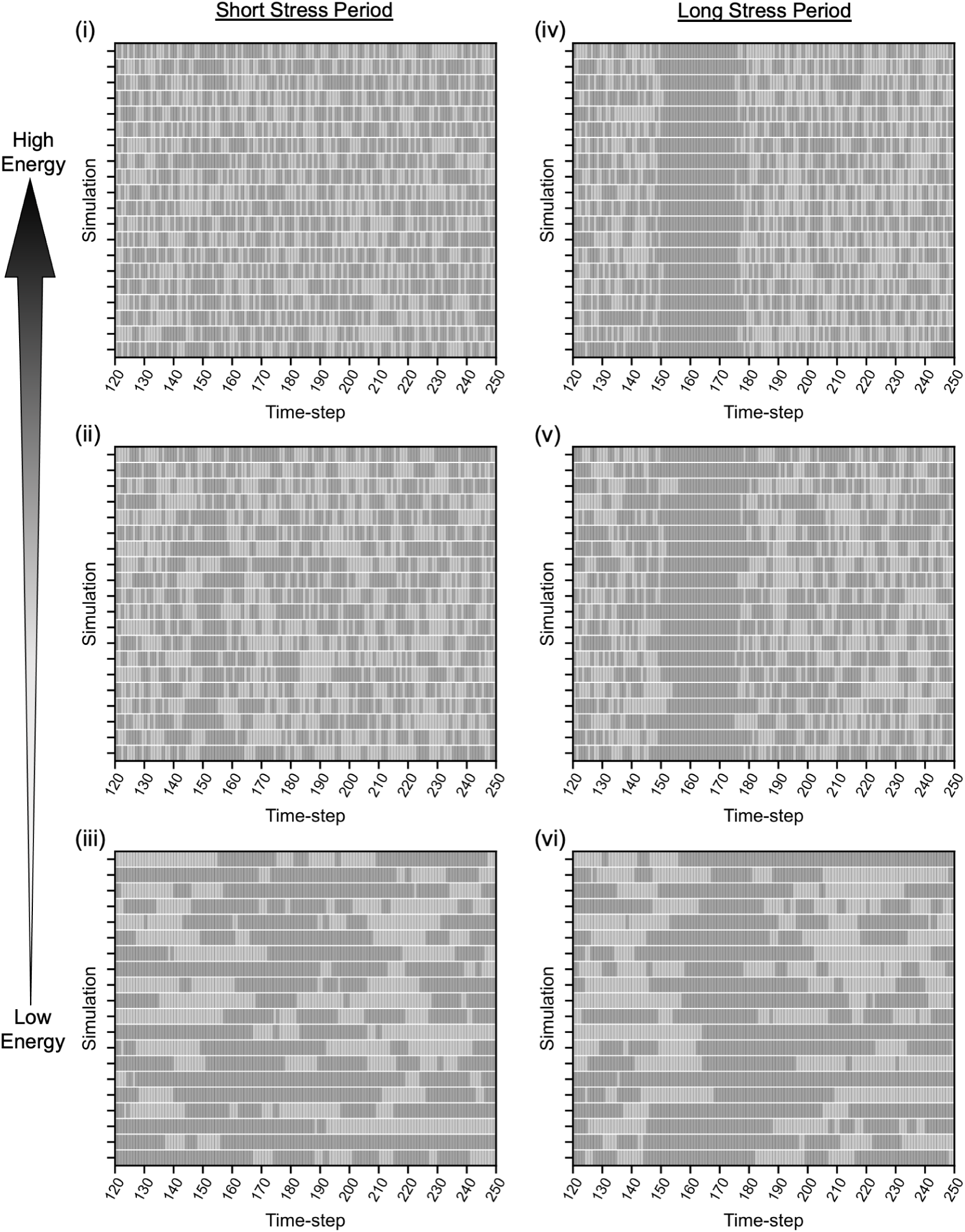
*E. coli* displays significant heterogeneity in *marRA* gene expression in the absence of stress. Qualitative plots displaying *marRA* “on”/“off” (dark/light grey) dynamics for 20 randomly selected individual *E. coli* simulations, when exposed to a (i)-(iii) short or (iv)-(vi) long stressor period. Panels display *marRA* dynamics between timesteps *t* = 120-250 at low, intermediate, and high energy availability. The short stressor period is a single timestep pulse activated at *t* = 150 and the long stressor period is activated at *t* = 150 until *t* = 174.

**Figure S7:**
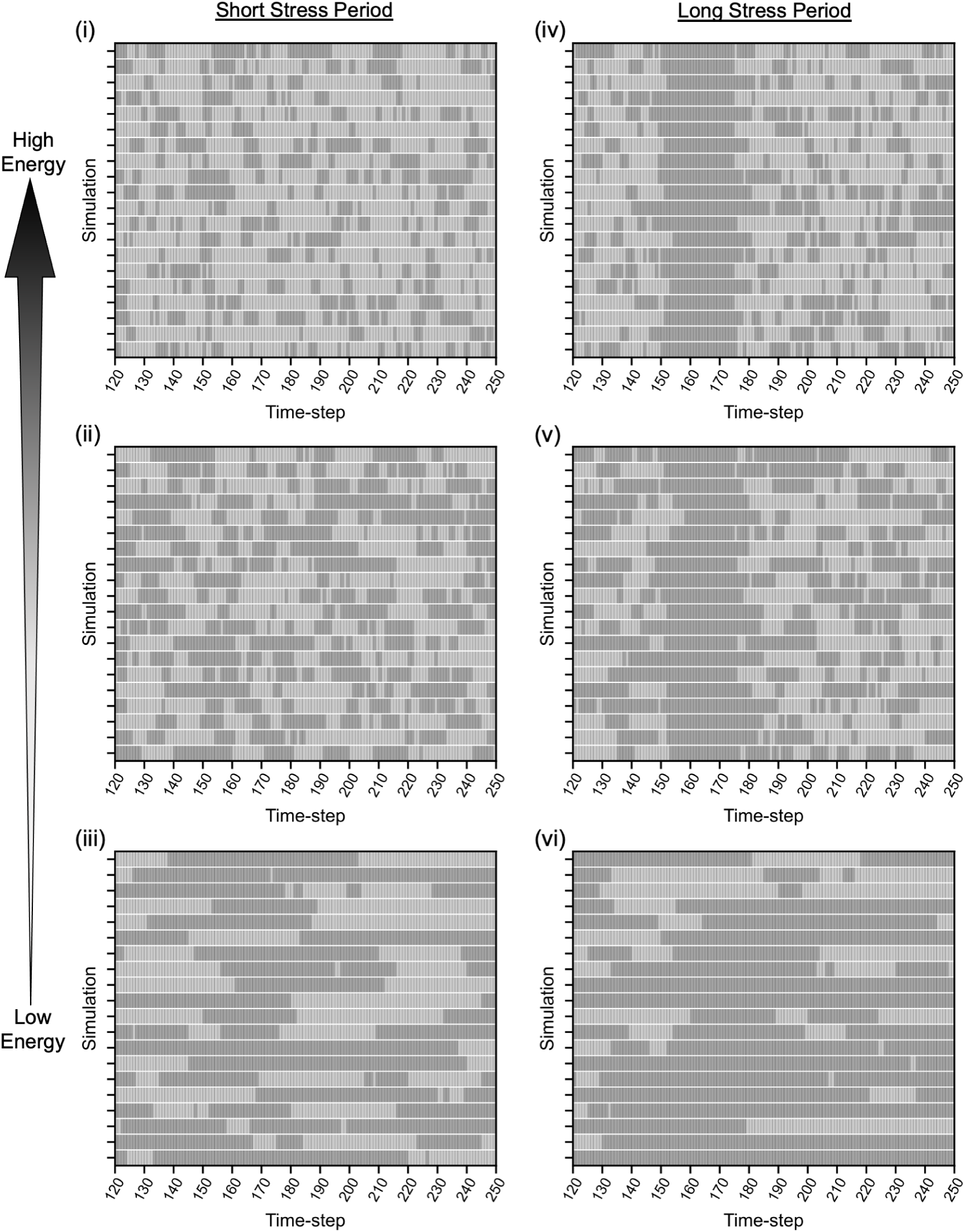
Simulations predict *Salmonella* expresses *acrAB* in heterogeneous pulses in the absence of stress. Qualitative plots displaying *acrAB* “on”/“off” (dark/light grey) dynamics for 20 randomly selected individual *Salmonella* simulations, when exposed to a (i)-(iii) short or (iv)-(vi) long stressor period. Panels display *acrAB* dynamics between timesteps *t* = 120-250 at low, intermediate, and high energy availability. The short stressor period is a single timestep pulse activated at *t* = 150 and the long stressor period is activated at *t* = 150 until *t* = 174.

**Figure S8:**
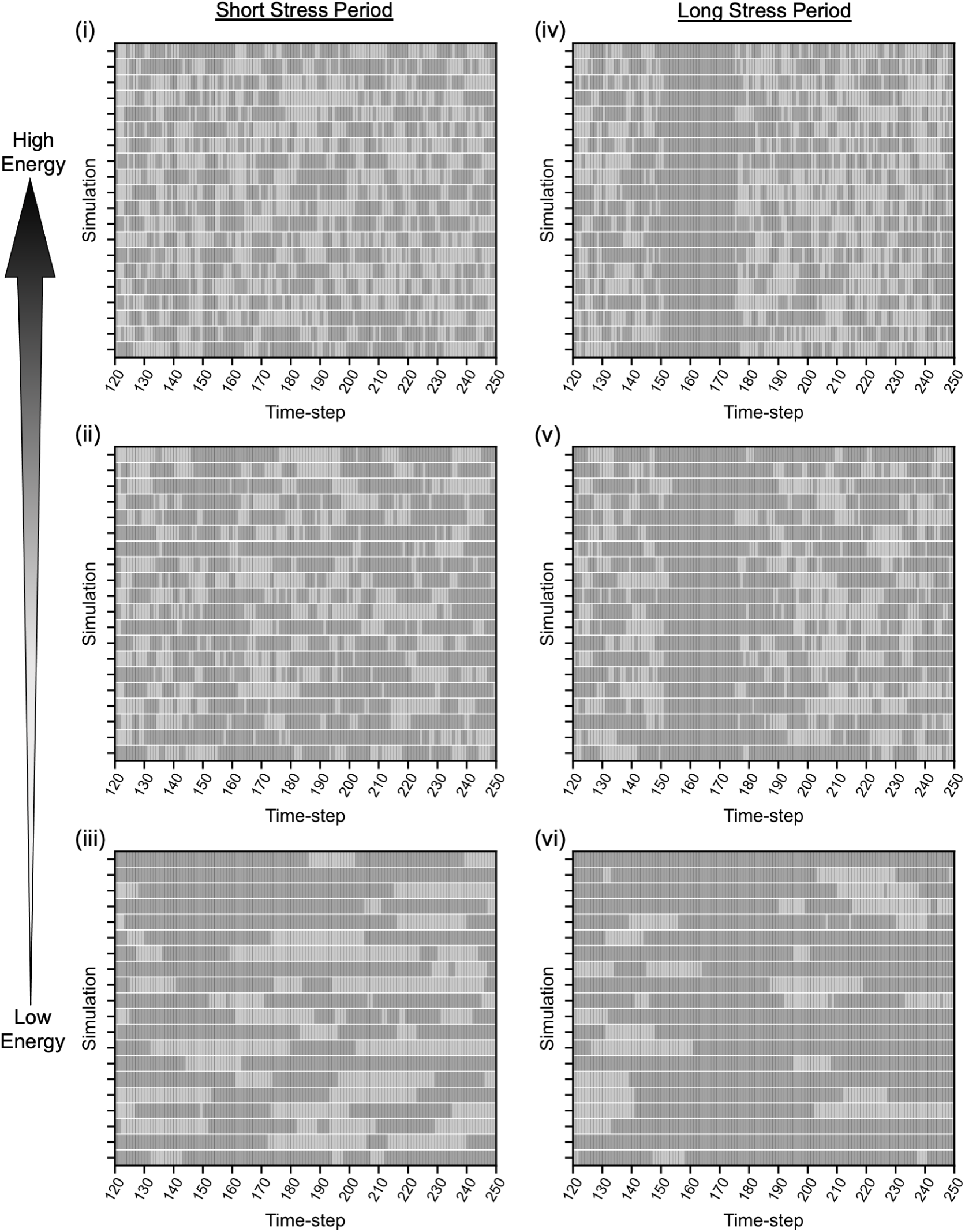
*Salmonella* displays significant heterogeneity in *ramA* gene expression in the absence of stress. Qualitative plots displaying *ramA* “on”/“off” (dark/light grey) dynamics for 20 randomly selected individual *Salmonella* simulations, when exposed to a (i)-(iii) short or (iv)-(vi) long stressor period. Panels display *ramA* dynamics between timesteps *t* = 120-250 at low, intermediate, and high energy availability. The short stressor period is a single timestep pulse activated at *t* = 150 and the long stressor period is activated at *t* = 150 until *t* = 174.

**Table S1:**
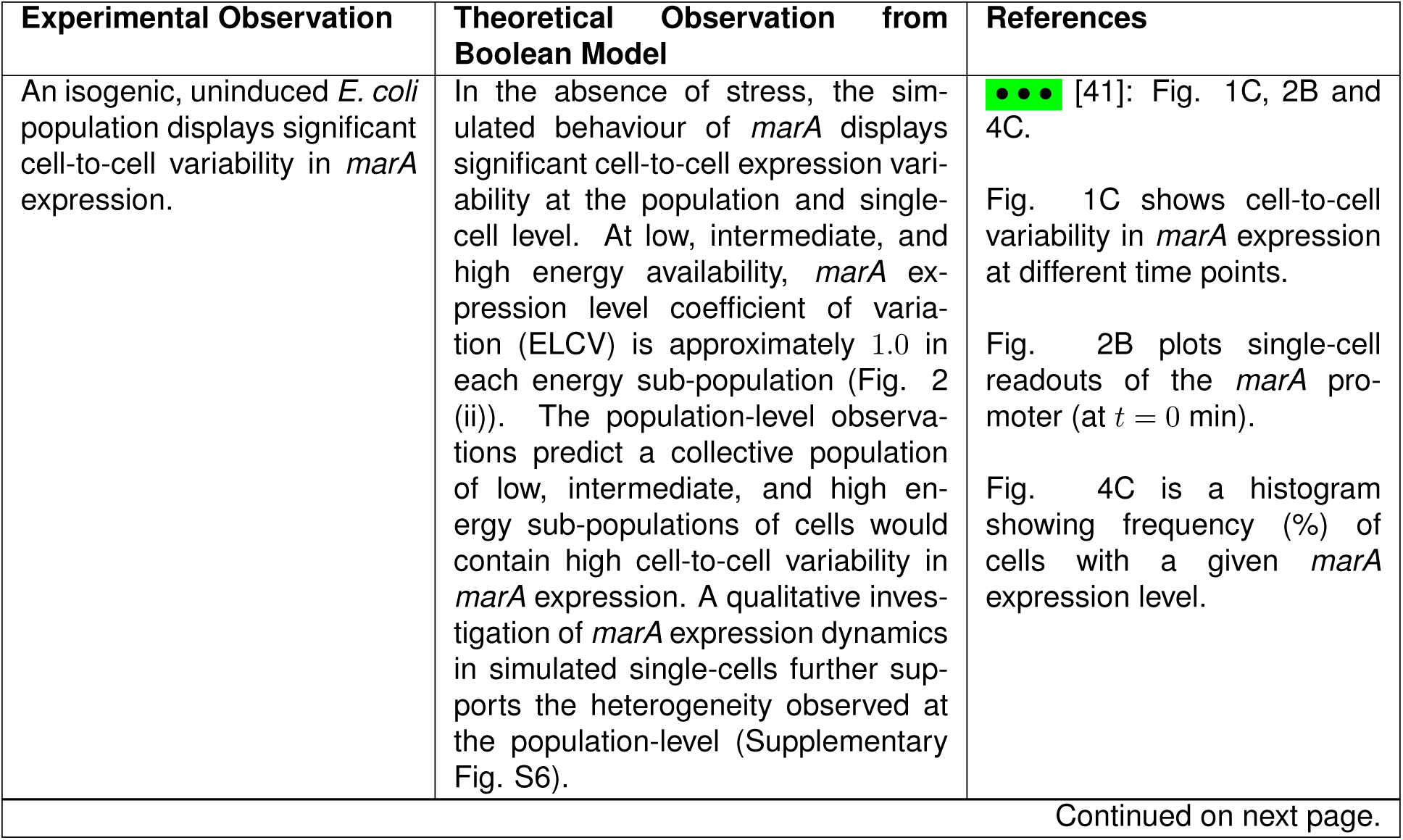
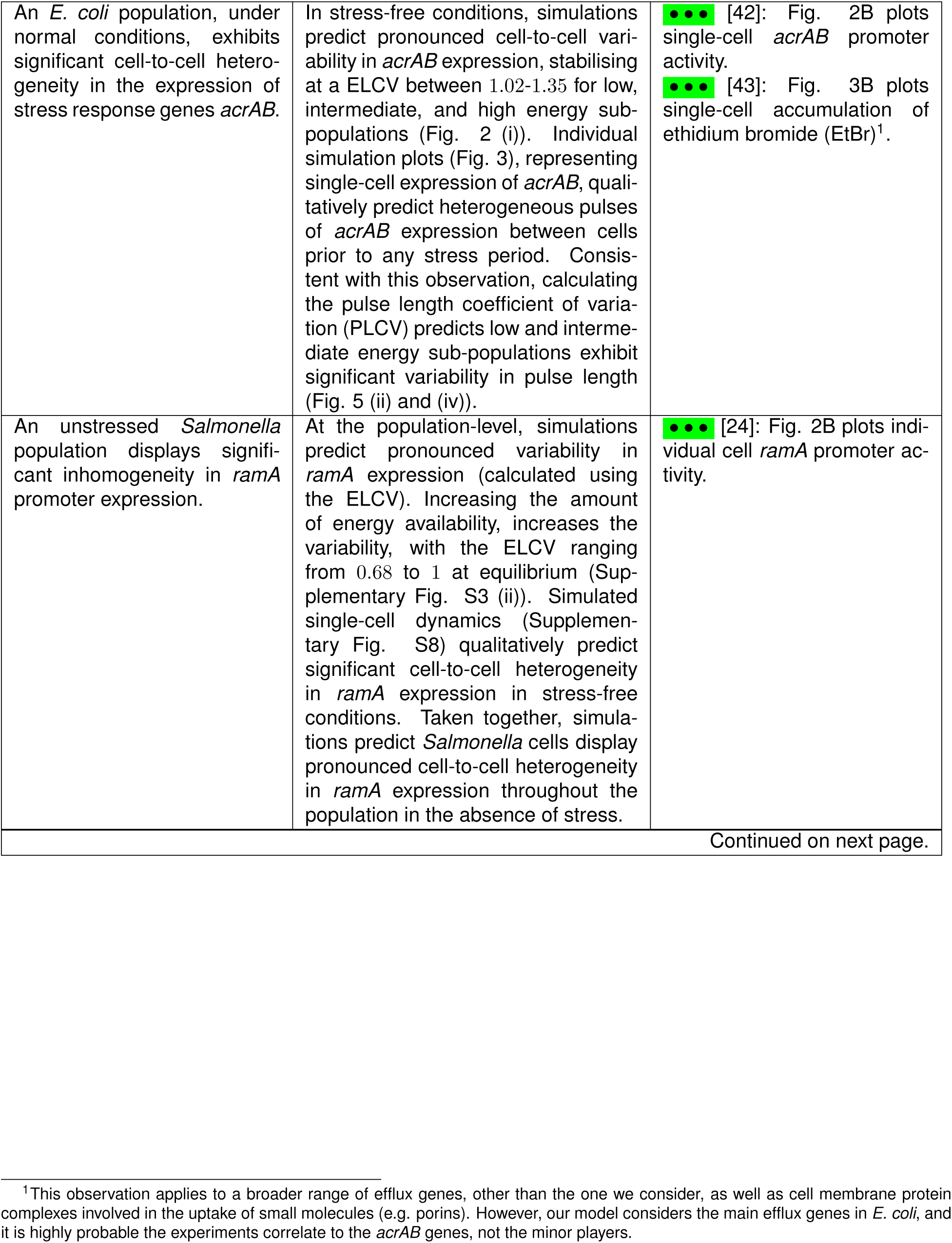
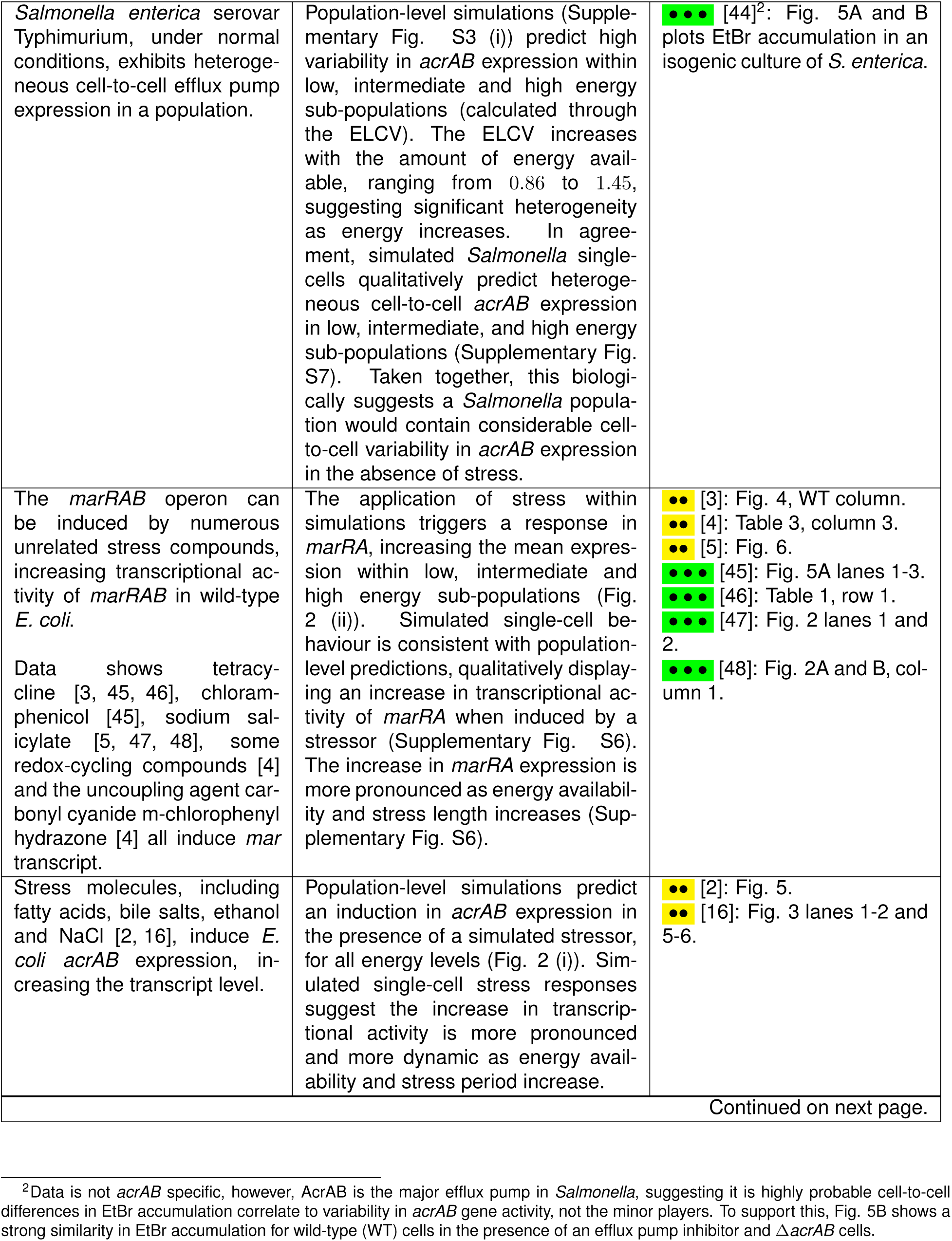
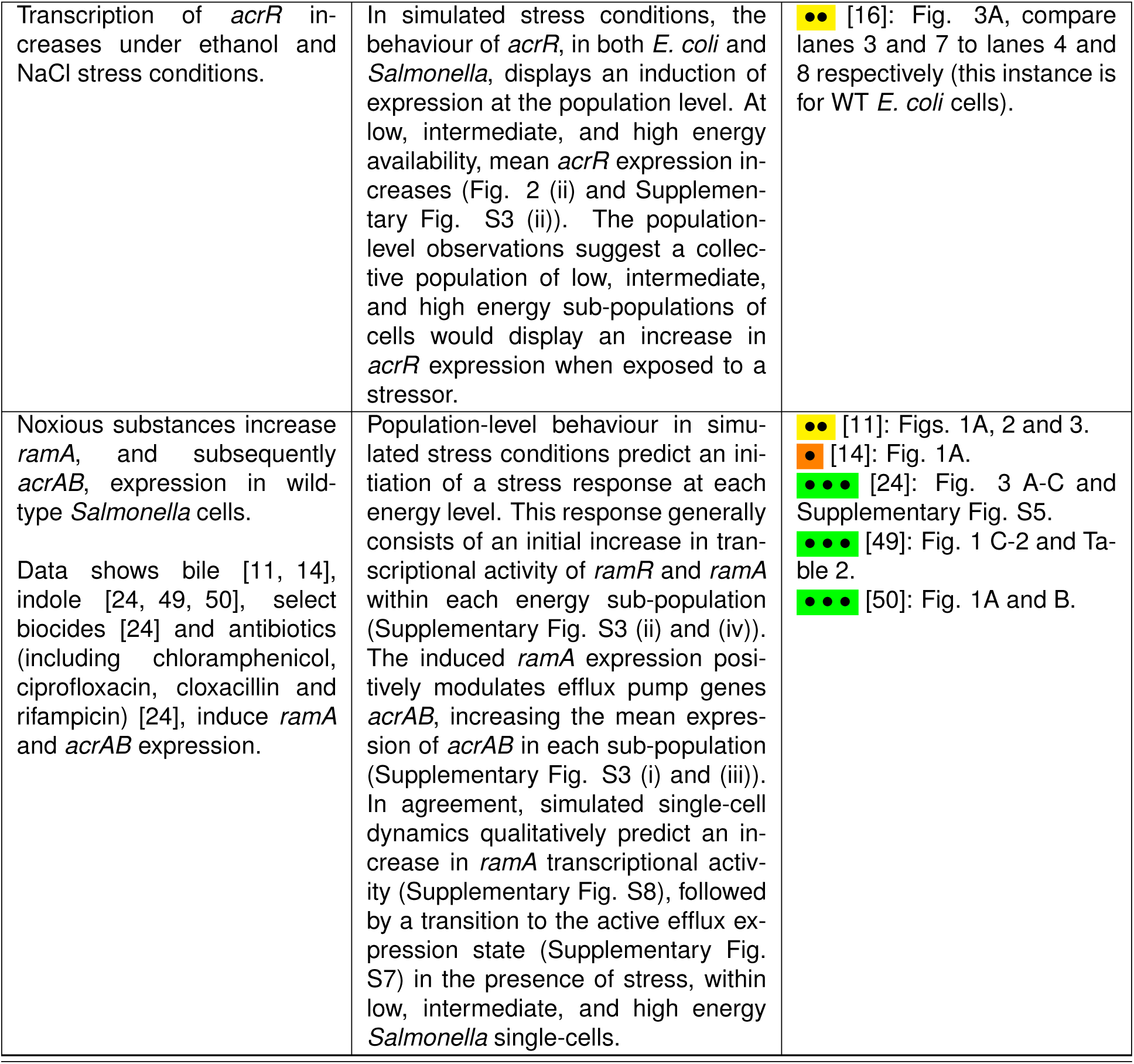
Theoretical model reproduces realistic behaviour in E. coli and Salmonella. Each row in the table describes experimentally observed behaviour (with references) and theoretical results arising from our model that capture analogous behaviour. References supporting experimental observations are labelled according to the dependence of information contained within each reference that is applied to construct the gene regulatory networks. The reference markers show: (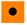) a dependent study where observations are used to flow into GRN construction, making them more difficult to disambiguate from model construction; (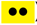) a study where the mechanism of the stated observation is unspecified, but other results are used in model construction; (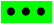) an independent study where the observation tests the model predictions. In summary, references with more dots have a higher strength of independence, and are a better validation of the model.

| Experimental Observation | Theoretical Observation from Boolean Model | References |
| --- | --- | --- |
| An isogenic, uninduced *E. coli* population displays significant cell-to-cell variability in *marA* expression. | In the absence of stress, the simulated behaviour of *marA* displays significant cell-to-cell expression variability at the population and single-cell level. At low, intermediate, and high energy availability, *marA* expression level coefficient of variation (ELCV) is approximately 1.0 in each energy sub-population (Fig. 2 (ii)). The population-level observations predict a collective population of low, intermediate, and high energy sub-populations of cells would contain high cell-to-cell variability in *marA* expression. A qualitative investigation of *marA* expression dynamics in simulated single-cells further supports the heterogeneity observed at the population-level (Supplementary Fig. S6). | ●●● [41]: Fig. 1C, 2B and 4C.<br>Fig. 1C shows cell-to-cell variability in *marA* expression at different time points.<br>Fig. 2B plots single-cell readouts of the *marA* promoter (at $t = 0$ min).<br>Fig. 4C is a histogram showing frequency (%) of cells with a given *marA* expression level. |
| Continued on next page. |  |  |
| An *E. coli* population, under normal conditions, exhibits significant cell-to-cell heterogeneity in the expression of stress response genes *acrAB*. | In stress-free conditions, simulations predict pronounced cell-to-cell variability in *acrAB* expression, stabilising at a ELCV between 1.02-1.35 for low, intermediate, and high energy sub-populations (Fig. 2 (i)). Individual simulation plots (Fig. 3), representing single-cell expression of *acrAB*, qualitatively predict heterogeneous pulses of *acrAB* expression between cells prior to any stress period. Consistent with this observation, calculating the pulse length coefficient of variation (PLCV) predicts low and intermediate energy sub-populations exhibit significant variability in pulse length (Fig. 5 (ii) and (iv)). | ●●● [42]: Fig. 2B plots single-cell *acrAB* promoter activity.<br>●●● [43]: Fig. 3B plots single-cell accumulation of ethidium bromide (EtBr)<sup>1</sup>. |
| An unstressed *Salmonella* population displays significant inhomogeneity in *ramA* promoter expression. | At the population-level, simulations predict pronounced variability in *ramA* expression (calculated using the ELCV). Increasing the amount of energy availability, increases the variability, with the ELCV ranging from 0.68 to 1 at equilibrium (Supplementary Fig. S3 (ii)). Simulated single-cell dynamics (Supplementary Fig. S8) qualitatively predict significant cell-to-cell heterogeneity in *ramA* expression in stress-free conditions. Taken together, simulations predict *Salmonella* cells display pronounced cell-to-cell heterogeneity in *ramA* expression throughout the population in the absence of stress. | ●●● [24]: Fig. 2B plots individual cell *ramA* promoter activity. |
| *Salmonella enterica* serovar Typhimurium, under normal conditions, exhibits heterogeneous cell-to-cell efflux pump expression in a population. | Population-level simulations (Supplementary Fig. S3 (i)) predict high variability in *acrAB* expression within low, intermediate and high energy sub-populations (calculated through the ELCV). The ELCV increases with the amount of energy available, ranging from 0.86 to 1.45, suggesting significant heterogeneity as energy increases. In agreement, simulated *Salmonella* single-cells qualitatively predict heterogeneous cell-to-cell *acrAB* expression in low, intermediate, and high energy sub-populations (Supplementary Fig. S7). Taken together, this biologically suggests a *Salmonella* population would contain considerable cell-to-cell variability in *acrAB* expression in the absence of stress. | ●●● [44]<sup>2</sup>: Fig. 5A and B plots EtBr accumulation in an isogenic culture of *S. enterica*. |
| The *marRAB* operon can be induced by numerous unrelated stress compounds, increasing transcriptional activity of *marRAB* in wild-type *E. coli*.<br>Data shows tetracycline [3, 45, 46], chloramphenicol [45], sodium salicylate [5, 47, 48], some redox-cycling compounds [4] and the uncoupling agent carbonyl cyanide m-chlorophenyl hydrazone [4] all induce *mar* transcript. | The application of stress within simulations triggers a response in *marRA*, increasing the mean expression within low, intermediate and high energy sub-populations (Fig. 2 (ii)). Simulated single-cell behaviour is consistent with population-level predictions, qualitatively displaying an increase in transcriptional activity of *marRA* when induced by a stressor (Supplementary Fig. S6). The increase in *marRA* expression is more pronounced as energy availability and stress length increases (Supplementary Fig. S6). | ●● [3]: Fig. 4, WT column.<br>●● [4]: Table 3, column 3.<br>●● [5]: Fig. 6.<br>●●● [45]: Fig. 5A lanes 1-3.<br>●●● [46]: Table 1, row 1.<br>●●● [47]: Fig. 2 lanes 1 and 2.<br>●●● [48]: Fig. 2A and B, column 1. |
| Stress molecules, including fatty acids, bile salts, ethanol and NaCl [2, 16], induce *E. coli* *acrAB* expression, increasing the transcript level. | Population-level simulations predict an induction in *acrAB* expression in the presence of a simulated stressor, for all energy levels (Fig. 2 (i)). Simulated single-cell stress responses suggest the increase in transcriptional activity is more pronounced and more dynamic as energy availability and stress period increase. | ●● [2]: Fig. 5.<br>●● [16]: Fig. 3 lanes 1-2 and 5-6. |
| Continued on next page. |  |  |

<sup>2</sup>Data is not *acrAB* specific, however, AcrAB is the major efflux pump in *Salmonella*, suggesting it is highly probable cell-to-cell differences in EtBr accumulation correlate to variability in *acrAB* gene activity, not the minor players. To support this, Fig. 5B shows a strong similarity in EtBr accumulation for wild-type (WT) cells in the presence of an efflux pump inhibitor and $\Delta$ *acrAB* cells.

|  |  |  |
| --- | --- | --- |
| Transcription of *acrR* increases under ethanol and NaCl stress conditions. | In simulated stress conditions, the behaviour of *acrR*, in both *E. coli* and *Salmonella*, displays an induction of expression at the population level. At low, intermediate, and high energy availability, mean *acrR* expression increases (Fig. 2 (ii) and Supplementary Fig. S3 (ii)). The population-level observations suggest a collective population of low, intermediate, and high energy sub-populations of cells would display an increase in *acrR* expression when exposed to a stressor. | ●● [16]: Fig. 3A, compare lanes 3 and 7 to lanes 4 and 8 respectively (this instance is for WT *E. coli* cells). |
| Noxious substances increase *ramA*, and subsequently *acrAB*, expression in wild-type *Salmonella* cells.<br>Data shows bile [11, 14], indole [24, 49, 50], select biocides [24] and antibiotics (including chloramphenicol, ciprofloxacin, cloxacillin and rifampicin) [24], induce *ramA* and *acrAB* expression. | Population-level behaviour in simulated stress conditions predict an initiation of a stress response at each energy level. This response generally consists of an initial increase in transcriptional activity of *ramR* and *ramA* within each energy sub-population (Supplementary Fig. S3 (ii) and (iv)). The induced *ramA* expression positively modulates efflux pump genes *acrAB*, increasing the mean expression of *acrAB* in each sub-population (Supplementary Fig. S3 (i) and (iii)). In agreement, simulated single-cell dynamics qualitatively predict an increase in *ramA* transcriptional activity (Supplementary Fig. S8), followed by a transition to the active efflux expression state (Supplementary Fig. S7) in the presence of stress, within low, intermediate, and high energy *Salmonella* single-cells. | ●● [11]: Figs. 1A, 2 and 3.<br>● [14]: Fig. 1A.<br>●●● [24]: Fig. 3 A-C and Supplementary Fig. S5.<br>●●● [49]: Fig. 1 C-2 and Table 2.<br>●●● [50]: Fig. 1A and B. |

## Notes

https://github.com/StochasticBiology/boolean-efflux

### Competing Interest Statement

The authors have declared no competing interest.

